# Inhibition of BRN2 in Melanoma Reverses Anoikis Resistance and Sensitizes Cells to Killing by Vemurafenib

**DOI:** 10.1101/2025.07.31.667908

**Authors:** Hannah M. Neuendorf, Xie He, Mark N. Adams, Khoa A. Tran, Aaron G. Smith, Paul V. Bernhardt, Craig M. Williams, Jacinta L. Simmons, Glen M. Boyle

## Abstract

Anoikis is an apoptotic cell death program triggered upon detachment from surrounding extracellular structures. However, the ability to evade cell death by anoikis in the presence of apoptosis-inducing stimuli is necessary for the formation of malignant tumors and progression to metastasis. Our findings indicate that the BRN2 *(POU3F2)* transcription factor is associated with anoikis resistance in melanoma cells. However, the BRN2 signaling cascade driving anoikis resistance remains unknown. Herein, we employed genome-wide CRISPR screens to validate BRN2 as a driver of anoikis resistance. Small molecule inhibition of BRN2 in melanoma cell lines with acquired anoikis resistance resensitized to death by anoikis in ultra-low attachment conditions. Our quantitative mass spectrometry analysis revealed that BRN2 functionally impacts oxidative phosphorylation and mitochondrial activity, whereby probes designed to inhibit BRN2 induced apoptosis and mitochondrial fragmentation through the MAPK and NF-κB signaling pathways and reduction in PPARɣ expression. Our study suggests that inhibition of BRN2 might allow the targeting of metastatic cells in circulation, and sensitizes cells to BRAF-targeted therapy, improving the prognosis for melanoma patients.

Graphical Abstract.
Role of BRN2 in driving anoikis resistance in melanoma. Upon detachment from the extra-cellular matrix (ECM) melanoma cells must evade cell death by anoikis to seed distant metastases. This study expanded the understanding of the role of the BRN2 transcription factor as a driver of resistance to anoikis in melanoma. The use of small molecule inhibitors targeting BRN2 revealed that the transcription factor drives anoikis resistance via the MAPK and NF-κB signaling pathways, resulting in PPARγ dysregulation and subsequently driving mitochondrial dysfunction. Green boxes = previously published drivers of anoikis resistance in melanoma. Blue box = changes to mitochondrial function following inhibition of BRN2 as determined by proteomics analysis.

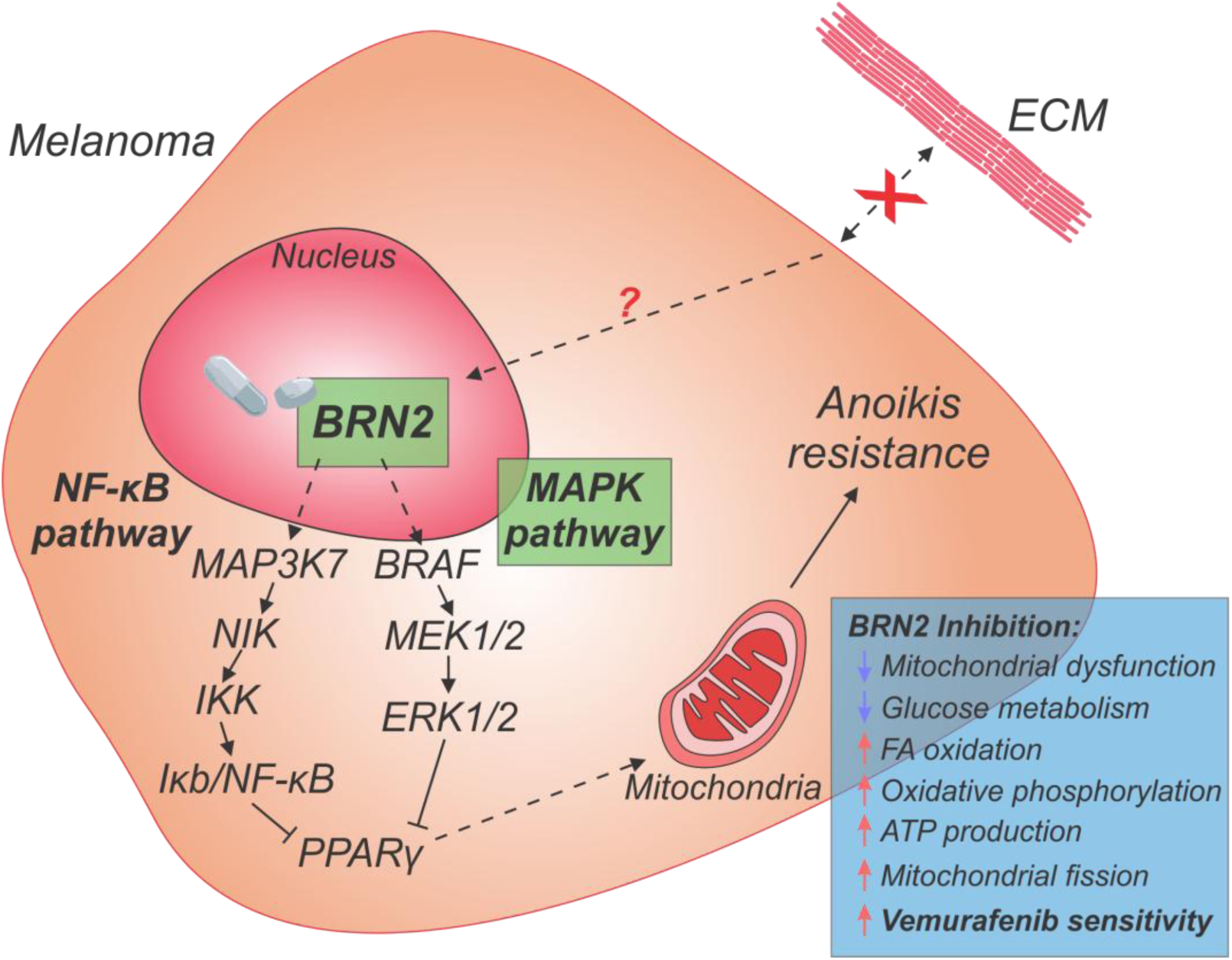

## 1.0 Introduction

Cutaneous Malignant Melanoma, a cancer derived from the oncogenic transformation of melanocytes, remains a significant public health burden [1]. This is particularly true in populations with exposure to high levels of ultra-violet radiation (UVR), which is one of the major risk factors for the development of melanoma resulting from photodamage of DNA by UVR [2, 3]. Early-stage melanoma is cured in the majority of cases through surgical resection of the primary tumor with a margin of surrounding healthy tissue [4]. However, a proportion of patients experience disease recurrence or are diagnosed with disseminated disease [5].

A range of therapies have been trialed and approved for the treatment of patients with advanced disease. The advent of MAPK pathway targeted inhibitors against BRAF and MEK [6–9], and immune checkpoint inhibitors (ICIs) [10–13], have resulted in an increase in the progression free survival of patients with stage IV metastatic melanoma to 31 % at 10 years, and overall survival of 43 %, with current best-practice treatment [14]. Despite these therapies, more than half of patients succumb to the disease due to both intrinsic and acquired therapy resistance [15]. Therefore, a greater understanding of the melanoma cell populations and biological mechanisms driving therapy resistance and recurrence are required to identify novel treatments and improve patient prognosis.

Anoikis, derived from the Ancient Greek ἄνοικος (ánoikos) meaning ‘homeless wanderer’, describes the apoptotic cell death program triggered in almost every cell type in the human body upon detachment from surrounding extracellular structures [16–18]. It is a physiological mechanism that prevents the growth of cells at inappropriate locations within the body, and maintains an equilibrium of cell number within tissues [19]. Mechanistically, anoikis is triggered through a combination of both intrinsic and extrinsic apoptosis pathways converging at the mitochondria and resulting in the activation of caspases, triggering DNA fragmentation and cell death [20]. As a hallmark of cancer, melanoma cells must develop mechanisms to resist anoikis to survive the process of disseminating into circulation to subsequently seed metastatic lesions at secondary sites [20]. While cancers of true epithelial origin are known to evade anoikis by hijacking mechanisms involved in epithelial-mesenchymal transition (EMT) [21], the origin of melanocytes in the neural crest means that they do not undergo a true-EMT [22, 23]. Consequently, only a few key molecules have been found to contribute to anoikis resistance in melanoma (reviewed in [24]). For instance, activation of the MAPK and PI3K-Akt pathways via integrin stimulation, as well as mutations in *BRAF, NRAS* or *NF1,* have been associated with increased resistance to cell death by anoikis [25–30].

BRN2, the POU domain transcription factor encoded by the gene *POU3F2,* is a neural-lineage transcription factor involved in the migration of neural crest cells and mediates subsequent melanocytic lineage development and differentiation [31–33]. While BRN2 is not expressed in melanocytes, it is upregulated in melanoma cells following *BRAF* mutation and p16 loss, and is associated with the acquisition of a highly invasive and motile phenotype that is thought to drive metastasis formation [34, 35]. High levels of BRN2 expression in melanoma cells are linked to a slow-cycling, dedifferentiated, stem cell-like phenotype [36–39], and the upregulation of invasive signature genes such as *MET, STAT3, TWIST, ITGB1* (encoding β1 integrin) and *PPARG* [40]. BRN2 is further implicated in downregulating the proliferative phenotype via reciprocal regulation of MITF transcription factor expression [37, 41–43].

In this study, we utilized genome-wide CRISPR screening technology to uncover drivers of anoikis resistance in melanoma and validated the role of BRN2 as a driver of anoikis resistance. We then employed two BRN2 small molecule inhibitors as probes to better understand the signaling network regulated by BRN2 that promotes anoikis resistance. In doing so, we identified the potential for BRN2 as a druggable target for the killing of melanoma cells under non-adherent culture conditions, particularly in combination with the BRAF inhibitor, vemurafenib. Identifying and targeting novel molecules that function in anoikis resistance with therapeutic strategies has the potential to prevent the dissemination of cells from the primary tumor and allow the targeting of metastatic cells in circulation, improving the prognosis for patients with melanoma.

## 2.0 Materials and Methods

### 2.1 Cell lines

Melanoma cell lines used in this study have been described previously [44, 45] and were grown in RPMI 1640 media containing phenol red (Invitrogen, Waltham, MA), supplemented with 10 % fetal bovine serum (FBS) (Thermo Fisher Scientific, Waltham, MA). HEK293FT cells (Thermo Fisher Scientific) were grown in DMEM supplemented with 0.01 % GlutaMax^TM^ (Thermo Fisher Scientific), 0.01 % 100 mM Sodium Pyruvate, 1× MEM Non-Essential Amino Acids (Thermo Fisher Scientific) and 10 % FBS (Thermo Fisher Scientific). Cells were maintained at 37 °C with 5 % CO₂ in a 95 % humidified atmosphere. Regular mycoplasma testing was performed, and results were negative. Cell line identity was verified by short tandem repeat (STR) profiling with the GenePrint® 10 System (Promega, Madison, WI) according to the manufacturer’s instructions.

### 2.2 Genome-wide pooled lentiviral CRISPR library screens

#### 2.2.1 Generation of stable dCas9 cell lines

The SP-dCas9-VPR plasmid for CRISPRa was a gift from George Church (Addgene plasmid #63798; http://n2t.net/addgene:63798; RRID:Addgene_63798) and the lenti_dCas9-KRAB-MeCP2 plasmid for CRISPRi was a gift from Andrea Califano (Addgene plasmid #122205; http://n2t.net/addgene:122205; RRID:Addgene_122205).

MM253, MM370 and MM576 cell lines were transfected with 20 µg SP-dCas9-VPR plasmid in a T75 flask using 40 µL X-tremeGENE HP DNA Transfection Reagent (Sigma-Aldrich) in 2 mL Opti-MEM (Thermo Fisher Scientific) (2:1 transfection reagent to DNA ratio). A no-plasmid control was included in all transfections. Cells were selected with 200 µg/mL G-418 (Thermo Fisher Scientific) for MM253, and 400 µg/mL for MM370 and MM576 48 h after transfection. The lenti_dCas9-KRAB-MeCP2 plasmid DNA was packaged into lentivirus using an optimized protocol for third-generation lentiviruses [46]. Plasmid DNA (10 µg) was combined with pLP-1, pLP-2 and pVSV-G plasmids at a 2:2:1:1 ratio in 3 mL Opti-MEM media with 90 µL Lipofectamine 2000 (Invitrogen) before addition to 30 % confluent HEK293FT cells. Media was changed 24 h after transfection, and viral supernatant harvested after a further 48 h by centrifugation in a sterile Amicon Ultra-15 centrifugal filter unit (Merck, Darmstadt, Germany) at 4000 rpm, 4 °C for 30 min. Lentiviral particles were introduced into the cell lines using 6 µg/mL polybrene (Sigma-Aldrich) and transduced cells selected with blasticidin at 2 µg/mL for MM253 and MM370 cells, or 3 µg/mL for MM576 cells.

Activity of the dCas9 was verified in the cell lines (0.5 × 10^6^ per well in 6 well plates) by fluorescent reporter assay prior to conducting the CRISPR screens **(Supplementary Figure S1).** SP-dCas9-VPR activity was verified by co-transfecting the reporter-gT1 plasmid which was a gift from George Church (Addgene plasmid #47320; http://n2t.net/addgene:47320; RRID:Addgene_47320), M-SP-sgRNA plasmid (a gift from George Church (Addgene plasmid #48671; http://n2t.net/addgene:48671; RRID:Addgene_48671) with the pLenti6/V5-DEST^TM^ Gateway^TM^ Vector, previously altered in the laboratory with Enhanced Green Fluorescent Protein (EGFP) inserted between the attR1 and attR2 sites by LR recombination (pLenti6-EGFP) (Invitrogen). Activity of the lenti_dCas9-KRAB-MeCP2 was verified by co-transfection of the pU6-sgGFP-NT1 (a gift from Stanley Qi & Jonathan Weissman (Addgene plasmid #46914; http://n2t.net/addgene:46914; RRID:Addgene_46914) and pMLS-SV40-EGFP (a gift from Stanley Qi & Jonathan Weissman (Addgene plasmid #46919; http://n2t.net/addgene:46919; RRID:Addgene_46919). Co-transfections were performed at a 6:1 sgRNA to reporter ratio using the X-tremeGENE HP DNA Transfection Reagent (Sigma-Aldrich). Cells were fixed in 4 % paraformaldehyde (PFA) for 20 min, stained with 30 µg/mL hoescht 33342 in 1× PBS and imaged on the EVOS FL Auto Microscope (Thermo Fisher Scientific). Fluorescence was quantitated using QuPath-0.2.3 software, and the percentage of cells expressing the fluorescent constructs in the dCas9-expressing lines compared to the parental cell lines.

#### 2.2.2 Single-guide RNA library preparation and viral titer

The protocol for preparing the CRISPR sgRNA libraries was adapted from the Weissman lab protocols (available at https://weissman.wi.mit.edu/resources/). The Human genome-wide CRISPRa-v2 and CRISPRi-v2 libraries were a gift from Jonathan Weissman (Addgene ID #83978; http://n2t.net/addgene:83978; RRID:Addgene_83978 and Addgene ID #83969; http://n2t.net/addgene:83969; RRID:Addgene_83969) [47]. The libraries were transformed into MegaX DH10B T1^R^ Electrocomp™ Cells (Thermo Fisher Scientific) by electroporation in Gene Pulser/MicroPulser Electroporation Cuvettes, 0.2 cm gap (Bio-Rad Laboratories, Hercules, CA), at 2.0 kV, 2000 Ohms, 25 µF using an exponential waveform. Bacteria were immediately recovered in 1 mL S.O.C. medium (Invitrogen), then incubated at 37 °C with shaking at 230 rpm for 2 h. Half the volume of each culture was serial diluted in S.O.C (two step: 1/50 then 1/100) and 50 µL of the final dilution spread on LB agar plates containing 100 µg/mL ampicillin. The other half was used to inoculate 500 mL LB broth with 100 µg/mL ampicillin. Both were incubated at 37 °C overnight. Colonies were counted on agar plates to calculate transformation efficiency, and DNA extracted from liquid cultures using the Qiagen Plasmid Mega kit per the manufacturer’s protocol. Purified plasmid DNA was packaged into lentivirus using eleven T75 flasks per library as described above, and viral titer (functional transducing units (TU/mL)) calculated per the Gill and Denham protocol [46] using flow cytometry to detect the presence of tagBFP in the hCRISPRa/i-v2 plasmid backbone. Using a multiplicity of infection (MOI) of 0.3, and allowing for sgRNA coverage of 100-fold, the number of cells and volume of virus necessary to maintain sgRNA enrichment for the subsequent CRISPR screens were calculated.

#### 2.2.3 CRISPR screening for drivers of anoikis resistance

To conduct the CRISPR screens, cell lines with stable dCas9 expression were transduced with the hCRISPRa-v2 or hCRISPRi-v2 sgRNA libraries as determined by viral titer experiments. After 48 h, transduced cells were selected with 5 µg/mL puromycin for 4 days. Viable cells (1 × 10^6^, per trypan blue staining) were snap frozen as a day 0 control, 2 × 10^6^ cells plated into a T75 flask as an adherent control, or cells plated into ultra-low attachment at 100-fold sgRNA enrichment. After 7 days, adherent controls were trypsinized, and pellets snap frozen. Viable cells from non-adherent cultures were collected by staining with 3 µM propidium iodide for 15 min at room temperature then FACS sorting on the BD FACS Aria^TM^ II (BD Biosciences), before snap freezing. Three independent biological replicates were performed for the three melanoma cell lines for both the CRISPRa and CRISPRi screens.

#### 2.2.4 Sample preparation, sequencing and analysis

DNA was extracted from defrosted pellets using the QiaAmp DNA Mini Kit (Qiagen) following the manufacturer’s protocol. The Weismann laboratory protocol for ‘Illumina sequencing sample preparation for use with CRISPRi/a-v2 libraries’ was used as a baseline methodology for the preparation of samples for sequencing. PCR was performed on each of the CRISPR samples to amplify the sgRNAs, and incorporate the 3’ and 5’ Illumina adapters and the index (barcode for de-multiplexing) per the protocol. Primers were ordered from Sigma-Aldrich with standard desalting. PCR products were purified using SPRIselect beads (Beckman Coulter, Brea, CA) on 150 µL DNA at three different ratios (0.65×, 1× then 0.9×) sequentially to remove genomic DNA and primer dimers. Quality control was performed using the D1000 TapeStation on 1 µL of DNA from each sample to confirm purity and concentration.

Samples were pooled and sequenced using 51 bp single-end reads with at least 50 million reads per sample on Illumina next-generation sequencing systems; NovaSeq6000 and NextSeq550 for CRISPRa, NextSeq2000 for CRISPRi. Sequencing primers were: 5’ Sequencing Primer for set ‘A’ samples GTGTGTTTTGAGACTATAAGTATCCCTTGGAGAACCACCTTGTTG, 3’ Sequencing Primer for set ‘B’ samples CCACTTTTTCAAGTTGATAACGGACTAGCCTTATTTAAACTTGCTATGCTGT. Raw files were de-multiplexed using the TruSeq Indexes, and converted to .fastq files using Bcl2fastq by the Sample Processing Facility at QIMR Berghofer.

Sequencing data was analyzed using the ScreenProcessing Analysis pipeline (https://github.com/mhorlbeck/ScreenProcessing/) as a template [47] and altered to suit the needs of the project. Full versions of the altered scripts for CRISPR analysis, and those used to generate graphs can be found at https://github.com/HannahNeuendorf/CRISPRAnalysis/. Statistical analyses and data manipulation were performed using the CRISPR_summary_stats.py script to calculate the number of genes from the library identified in the screen samples, and those missing. Volcano plots were generated from the log2 fold-change and Mann-Whitney p-values using volcanoplots.py script. Dependencies for the python analysis scripts are summarized in **Table 1**.

**Table 1.** Dependencies for CRISPR screen analysis scripts in Python.

| Script | Spyder environment | Python version | Package | Package version |
| --- | --- | --- | --- | --- |
| Fastqgz_to_counts_working.py | V5.0.5 | V3.6 |  |  |
| Process_experiments_working.py | V5.1.5 | V3.9 | Pandas<br>Scipy<br>Matplotlib | V0.23.0<br>V0.19<br>V2.2.2 |
| CRISPR_summary_stats.py | V5.1.5 | V3.9 | Numpy | V1.23.3 |
|  |  |  | Pandas<br>plotly | V1.4.4<br>V5.10.0 |
| Volcanoplots.py | V5.1.5 | V3.9 | Numpy<br>Pandas<br>plotly | V1.23.3<br>V1.4.4<br>V5.10.0 |

Significantly altered genes were analyzed using the QIAGEN Ingenuity Pathway Analysis (IPA) software and Pre-ranked Gene Set Enrichment Analysis (GSEA) v4.3.2 software (Broad Institute). Gene sets used for analysis were the hallmark (H1), curated (C2), ontology (C5) and oncogenic (C6) gene sets. Normalized enrichment score (NES) and nominal p-value were used to identify significantly enriched pathways within each gene set.

### 2.3 Anoikis assay

Melanoma cell lines (1 × 10^6^) were seeded in an ultra-low attachment 6-well plates (Corning, Corning, NY) in 2 mL media and allowed to grow for 7 days. For anoikis assays with BRN2 inhibitor treatment, 20 µM of B18, B18-94 or an equivalent volume of vehicle (DMSO) were added on days 0 and 4 of culture. Cells were stained on day 7 with 3 µM calcein-AM (Invitrogen) for screening of intrinsic anoikis resistance, or 3 µM CellTrace^TM^ Calcein Violet-AM (Thermo Fisher Scientific) for later experiments, plus 3 µM propidium iodide (Sigma-Aldrich, St Louis, MO) in 1× PBS for 15 min at room temperature, protected from light. Percentage viability was quantitated on the BD LSRFortessa^TM^ 4 laser cytometer (BD Biosciences, Franklin Lakes, NJ) using the 561 nm laser with the YG 670/30 filter to detect the PI-stained dead cells, and the 488 nm laser with the B 530/30 filter to detect the calcein-AM stained viable cells, or V 450/50 filter for calcein violet-AM stained viable cells. Live and dead cell populations were consistently gated using the BD FlowJo^TM^ Software v10.6.1, and viability calculated as a percentage of the total population of cells. Cell lines with viability below 20 % were designated ‘anoikis sensitive’, while those with viability above 60 % were termed ‘anoikis resistant’.

### 2.4 Western blot analysis

To generate samples, MM383, C-32 and MM386 cell lines were treated in adherent culture with vehicle, B18 or B18-94 at 20 µM for 3 or 7 days. Western blot analysis has been described previously [48]. Antibodies used in this study were: Brn2/POU3F2 (D2C1L) Rabbit mAb #12137 (CST) (1:1000), MITF (D5G7V) Rabbit mAb #12590 (CST) (1:1000), GAPDH MAB5718 (R&D Systems, Minnesota) (1:8000).

### 2.5 BRN2 inhibitor synthesis

The BRN2 small molecule inhibitors B18 and B18-94 were synthesized based on previously described methods by Thaper *et al.* [49]. Synthetic procedures, full characterization, x-ray crystallography, and copies of ^1^H and ^13^C nuclear magnetic resonance spectra can be found in **Supplementary File S1.**

### 2.6 Determination of BRN2 Inhibitor IC_50_

Cells were plated into 96 well plates (Corning) at 5000 cells per well for MM383, and 7500 cells per well for MM386 and C-32 cell lines. After 24 h, inhibitors or DMSO, were added at 10-fold serial dilutions from 100 µM to 0.0001 µM in triplicate. Media without inhibitor or DMSO was also added to cells as a negative control. An SRB assay was performed after 7 days as described previously [41] to quantitate cell survival. Percentage cell survival relative to the negative control was calculated in Microsoft Excel, and IC_50_ calculated with a variable slope non-linear fit model for inhibitor vs. normalized response in GraphPad Prism v8.4.3.

### 2.7 BRN2 immunofluorescence

To examine BRN2 chromatin binding in cells treated with the BRN2 inhibitors, immunofluorescence was performed. MM383, MM386 and C-32 cells were plated into Cellvis 96 well glass-bottom plates at 2 × 10^4^ cells per well. Cells were allowed to adhere overnight then 20 µM B18, B18-94 or vehicle added to cells (12 technical replicates for B18 and B18-94, 8 for vehicle, 3 independent biological replicates). After 96 h, half of each treatment type was extracted with chilled pre-extraction buffer (10 mM HEPES (pH 8), 20 mM NaCl, 5 mM MgCl_2,_ 0.5 % Igepal CA-630) for 4 min on ice and then rinsed with 1× PBS. All wells were then fixed with 4 % paraformaldehyde for 20 min at room temperature. Samples were permeabilized with 0.5 % Triton X-100 in 1× PBS for 5 min then blocked in FBT buffer (5 % FBS, 1 % BSA, 0.05 % Tween-20, 10 mM Tris (pH 6.8), 100 mM MgCl_2_) for 1 h on ice. Primary Brn2/POU3F2 (D2C1L) Rabbit mAb #12137 (CST) was diluted 1:1000 in FBT buffer and cells incubated in antibody solution at 4 °C overnight in a humidified container. Secondary Alexa Fluor® 488 donkey anti-rabbit (1:300) was diluted in FBT buffer and incubated for 2 h at ambient temperature protected from light. Nuclei were stained with 30 µg/mL hoescht 33342 in 1× PBS for 20 min at room temperature, then cells imaged in 1× PBS. BRN2 imaging was performed using high content microscopy as previously described [50, 51] using an InCell Analyzer 6500 high content microscopy system (GE Healthcare Life Sciences, Australia).

Image analysis was performed using the CellProfiler software v3.1.9, with BRN2 staining intensity calculated and reported per nuclei within each field of view. A minimum of 1100 nuclei were analyzed per condition and replicate. Plots show BRN2 staining intensity between extracted and un-extracted populations within replicates.

### 2.8 Reattach and grow assay following inhibitor treatment

Cells (5 × 10^4^) were seeded in an ultra-low attachment 96-well plate (Corning) and drug treatment (B18 or B18-94) or vehicle (DMSO) added on days 0 and 4. For experiments combining BRN2 inhibitors with vemurafenib, 2 × 10^4^ cells were seeded. After 7 days, the total volume from each well was transferred to a standard adherent 96-well plate (Corning), with 100 µL fresh media added without inhibitors. After 3 days, viability was determined by fixing plates with methylated spirits and cell density determined by sulforhodamine B (SRB) assay as previously described [41]. The percentage of cells able to reattach and grow was compared to vehicle control using Microsoft Excel.

### 2.9 Apoptosis assay

Cells (1 × 10^6^) were seeded into ultra-low attachment 6-well plates (Corning) with B18, B18-94 or DMSO added on days 0 and 4. After 1, 3 and 7 days, 1 × 10^5^ cells were harvested and stained with FITC-Annexin V (BD Biosciences) and 3 µM propidium iodide (PI) per the manufacturers’ protocol in 1× Binding Buffer (BD Biosciences). Data was acquired on the BD LSRFortessa^TM^ 4 laser cytometer (BD Biosciences) and cell populations consistently gated using the BD FlowJo^TM^ Software v10.6.1. Three independent replicates were performed for each cell line, with replicates averaged.

### 2.10 Proteomics analysis

Cells were seeded into T75 flasks (Falcon) at 15-20 % confluency with B18, B18-94 or DMSO added after 24 h and 4 days. After 7 days, cell pellets were collected from three independent biological replicates for quantification of whole proteome expression using Liquid Chromatography-Mass Spectrometry (LC-MS) by the QIMR Berghofer Proteomics Facility. Cell pellets were lysed in 100 mM triethylammonium bicarbonate (TEABC) with 4 % sodium deoxycholate (SDC) at 95 °C on a thermomixer for 5 min, followed by probe sonication. After removal of cell debris, protein concentration was quantified and normalized using a BCA Assay (Thermo Fisher Scientific). Protein was reduced and alkylated by addition of 10 mM tris(2-carboxyethyl)phosphine (TCEP) and 40 mM chloroacetamide (CAA) and incubated at 45 °C for 10 min. Samples were digested overnight at 37 °C with Pierce MS grade trypsin (Thermo Fisher Scientific) at a protease:protein ratio of 1:100 (w/w). Samples were acidified with TFA and cleaned using styrenedivinylbenzene reverse phase sulfonate (SDB-RPS) material tips (Empore, Sigma-Aldrich). Cleaned peptides were dried on a vacuum centrifuge and reconstituted in mass spectrometry (MS) loading solvent (0.1 % formic acid (FA) in 2 % ACN). Concentration of samples were determined by Nanodrop and normalized.

LC-MS runs were performed on the Thermo Exactive HF-X mass spectrometer paired with a Thermo neoVanquish LC equipped with Thermo Acclaim PepMap100 trap (5 mm × 300 µm ID) and analytical (150 mm × 300 µm ID) columns. After column equilibration, peptides were resolved using a gradient starting at 6 % solvent B (0.1 % FA in 80 % ACN), increasing to 27 % solvent B over 32 min, 35 % solvent B over 12.5 min, then to 44 % solvent B over 5 min, at a flow rate of 1.5 µL/min. This was followed by column washing in 99.9 % solvent B. Solvent A consisted of 0.1 % FA in MS-grade water.

For each sample, 1 µg of peptide was injected and mass spectrometry data was acquired in positive ion mode with a DIA acquisition method using the parameters specified by [52]. Briefly, MS1 scans were acquired from 390-1010 m/z (mass-to-charge ratio) at a resolution of 60,000 with an Automatic Gain Control (AGC) target of 1e6 and a maximum injection time of 60 ms. MS2 data was acquired using a 16 m/z staggered window scheme at a resolution of 15,000. The AGC was set to 1e6 with a maximum injection time of 20 ms. Loop counts were set to 76 and normalized collision energy was set to 27.

Raw data was searched against the reviewed Uniprot human database (downloaded June 11, 2024), along with a list of common contaminants, using the FASTA digest library-free option in DIA-NN v1.9 [53] with default settings. The main report table from the DIA-NN output was filtered for q-value < 0.01 at the precursor and protein group level. The data was processed using the Tidyproteomics R package v1.8.1 [54]. Briefly, the quantitative peptide data was converted into protein data, which was then normalized using the ‘randomforest’ method. Differential expression analysis was performed using the ‘limma’ method within the Tidyproteomics package and adjusted p-value calculated using Benjamini-Hochberg procedure.

To further analyze the significantly enriched pathways, comparisons were made between proteomics and phospho-proteomics using Ingenuity Pathway Analysis (IPA) software (Qiagen). Gene identification, log_2_ fold-change and p-value were used to rank gene enrichment. Genes with a fold-change between -1 and 1 were excluded from analysis. Overlap between cell lines and treatments for canonical pathways and upstream regulators output by IPA and overlap with existing datasets were generated using Interactivenn.net.

### 2.11 MitoTracker Red FM assay

To examine changes in mitochondrial activity following BRN2 inhibition, a MitoTracker Red FM assay was performed. For the 3 day assay, MM383, C-32 and MM386 cells were seeded into 96 well plates at 2 × 10^4^ cells per well in triplicate and allowed to adhere for 24 h. For the 7 day assay, MM383 cells were seeded into 96 well plates at 5 × 10^3^ cells per well, and the MM386 and C-32 cell lines seeded at 7.5 × 10^3^ cells per well in triplicate. Cells were treated with 20 µM B18, 20 µM B18-94 or vehicle on days 0 and 4 of culture.

After the treatment period, mitochondria were stained with 200 nM MitoTracker Red FM (Invitrogen) in warm media for 45 min at 37 °C. Cell nuclei were then stained with 5 µg/mL hoescht 33342 in media for 20 min at 37 °C. Hoescht 33342 and MitoTracker Red positive cells were imaged on the Evos FL microscope (Invitrogen) and fluorescence quantitated using the Biotek Synergy Neo2 Plate Reader (Agilent). Three independent biological replicates were performed for both the 3 and 7 day experiments. Technical replicates were averaged, and then positive MitoTracker signal normalized to hoescht 33342 to account for differences in cell number. Fold-change in mitochondrial activity was calculated between BRN2 inhibitor and DMSO treated wells in Microsoft Excel. Unpaired t-tests with Welch’s correction were performed using GraphPad Prism v8.4.3.

### 2.12 Statistical analysis

Data is expressed as mean ± error (SD or SEM as indicated). All statistical significance was determined by GraphPad Prism 8.4.3 for Windows (GraphPad Software, San Diego, California USA, www.graphpad.com) using tests as indicated. *p* < 0.05 was considered statistically significant. * *p* < 0.05, ** *p* < 0.01, *** *p* < 0.001, **** *p* < 0.0001.

## 3.0 Results

### Genome-wide CRISPR screening identifies downregulation of MITF as a mechanism of anoikis resistance

With the aim to identify genes contributing to resistance to cell death by anoikis in melanoma, genome-wide pooled lentiviral CRISPR library screens were conducted **(Figure 1A)**. A panel of human metastatic melanoma cell lines were evaluated for their intrinsic sensitivity to anoikis to select cell lines for the screen **(Figure 1B)**. Cells were plated into ultra-low attachment conditions for 7 days, and percentage viability established by flow cytometry following staining with propidium iodide (PI) and calcein-AM. Those with a viability greater than 60 % were denoted anoikis resistant, while those with viability lower than 20 % were considered anoikis sensitive. Three anoikis sensitive cell lines were chosen for the CRISPR screens (MM253, MM370 and MM576) to ensure that cell survival was limited to mechanisms driven primarily by transcriptional alterations induced by the CRISPR libraries, rather than intrinsic survival mechanisms. Cell lines were subsequently engineered to stably express the appropriate dCas9 molecules for CRISPR activation (CRISPRa) and CRISPR inhibition (CRISPRi) screens, and dCas9 activity verified by fluorescent reporter assays **(Supplementary Figure S1A/B)**.

**Figure 1.**
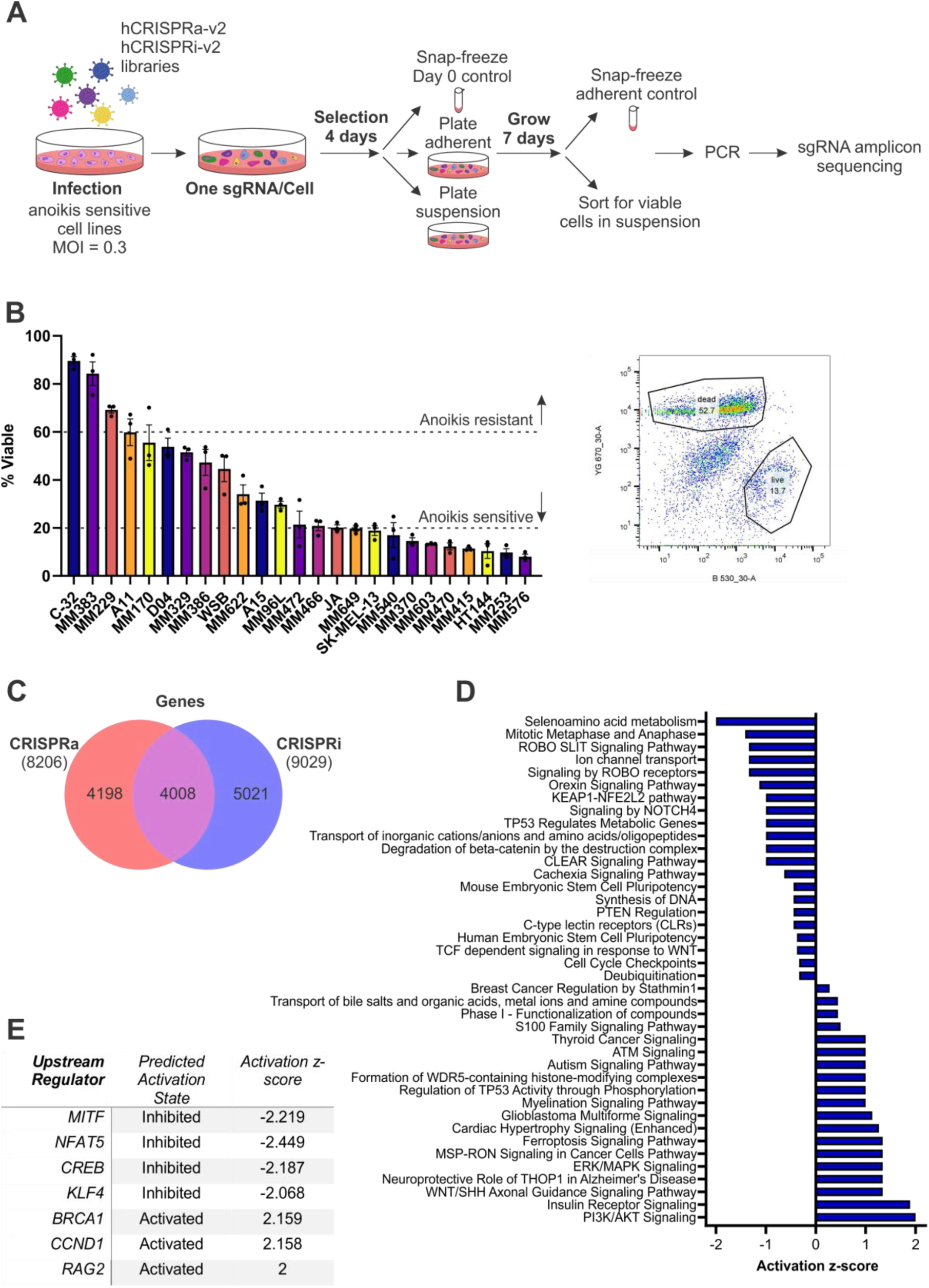
Genome-wide CRISPR screening for drivers of anoikis resistance. (A) Workflow for CRISPR activation (CRISPRa) and inhibition (CRISPRi) screens. Anoikis sensitive cell lines stably expressing dCas9 were transduced with either hCRISPRa-v2 or hCRISPRi-v2 sgRNA libraries followed by antibiotic selection. Cells were then snap-frozen as a Day 0 control, or plated under adherent or suspension culture conditions and allowed to grow for 7 days. Adherent control cells were then snap-frozen, and viable cells from the non-adherent culture collected by FACS using propidium iodide (PI) staining. Following PCR to incorporate barcodes for Illumina sequencing, PCR products were purified, and samples sequenced in a high-throughput manner. Enrichment of sgRNA in each sample was quantified and comparisons drawn between conditions using a python analysis pipeline. (B) Determination of baseline anoikis resistance. A panel of human melanoma cell lines were screened for their intrinsic sensitivity to anoikis. Cells were grown in ultra-low attachment plates for 7 days then stained with PI and calcein-AM, and percentage viability assessed by flow cytometry. Cell lines with viability > 60 % were considered anoikis resistant; those with viability < 20 % anoikis sensitive. MM253, MM370 and MM576 cell lines were then selected as the anoikis sensitive cell lines for the CRISPR screen. Values indicate mean ± SEM, n = 3 independent biological replicates. (C) Sequencing data from the CRISPR screens (n = 3 independent biological replicates for 3 cell lines) were analyzed using a python analysis pipeline adapted from Horlbeck *et al.* 2016 [47]. Dysregulated genes (p-value < 0.05) were exported into excel for analysis. 4008 genes were found to be significantly dysregulated between both the CRISRa and CRISPRi screens. (D) Ingenuity pathway analysis (IPA) of the CRISPR screens. The screens identified a range of pathways dysregulated by ultra-low adherent culture conditions. (E) Analysis of upstream regulators by IPA predicted inhibition of the MITF transcription factor as a contributor to the anoikis resistant phenotype.

To conduct the screen, dCas9 expressing cell lines were transduced with either the hCRISPRa-v2 or hCRISPRi-v2 [47] top-5 library of sgRNA, and were then snap frozen at day 0 (baseline control), or plated into standard adherent or ultra-low attachment conditions for 7 days. Next generation sequencing analysis of viable cells demonstrated that a high level of library coverage was maintained throughout the CRISPR screens **(Supplementary Figure S1C)**. Overall, more than 4000 genes were found to be significantly altered between control and non-adherent samples in both the CRISPRa and CRISPRi screens **(Figure 1C, Supplementary Figure S1D/E, Supplementary File S2/S3)**. Dysregulated gene and pathway analysis using Ingenuity Pathway Analysis (IPA) software revealed a range of canonical pathways that were significantly enriched in cells surviving in detached growth conditions **(Figure 1D, Supplementary File S4)**. These analyses identified known mechanisms of anoikis resistance such as activation of PI3K/AKT and ERK/MAPK signaling, as well as a range of novel pathways. IPA also identified *MITF* transcription factor inhibition as an upstream driver of the anoikis resistant phenotype in these cell lines (activation z-score -2.219) **(Figure 1E)**.

### Small molecule inhibitors bind BRN2 in melanoma cells

High BRN2 expression contributes to cell survival under non-adherent conditions [40]. Given the reciprocal relationship between MITF and BRN2 [32, 42, 43, 55, 56], the downregulation of MITF in the CRISPR screens was consistent with an upregulation of BRN2 in anoikis resistance, validating our previous findings. While BRN2 has been identified to contribute to anoikis resistance, its role in the process remains unclear. We therefore sought to further investigate the role of BRN2 in anoikis resistance using the inhibitors described by Thaper *et al.* as probes [49].

To examine the effect of the BRN2 inhibitors in melanoma, B18 and its derivative B18-94 **(Figure 2A)** were applied to melanoma cell lines at concentrations determined by Thaper *et al.* **(Supplementary Figure S2)**. Western blotting for expression of BRN2 and MITF following treatment with 20 µM of the inhibitors for 3 or 7 days revealed a reduction in BRN2 expression compared to vehicle over time in both the C-32 and MM386 cell lines **(Figure 2B)**. While BRN2 inhibition was not expected to directly affect the level of BRN2 expression, this may indicate destabilization of the BRN2 protein when not bound to DNA and/or the blocking of a BRN2-BRN2 positive feedback loop. Furthermore, expression of MITF was unchanged following 3 days of treatment, with variations in MITF post-translational modification demonstrated in the C-32 and MM386 cell lines at 7 days.

**Figure 2.**
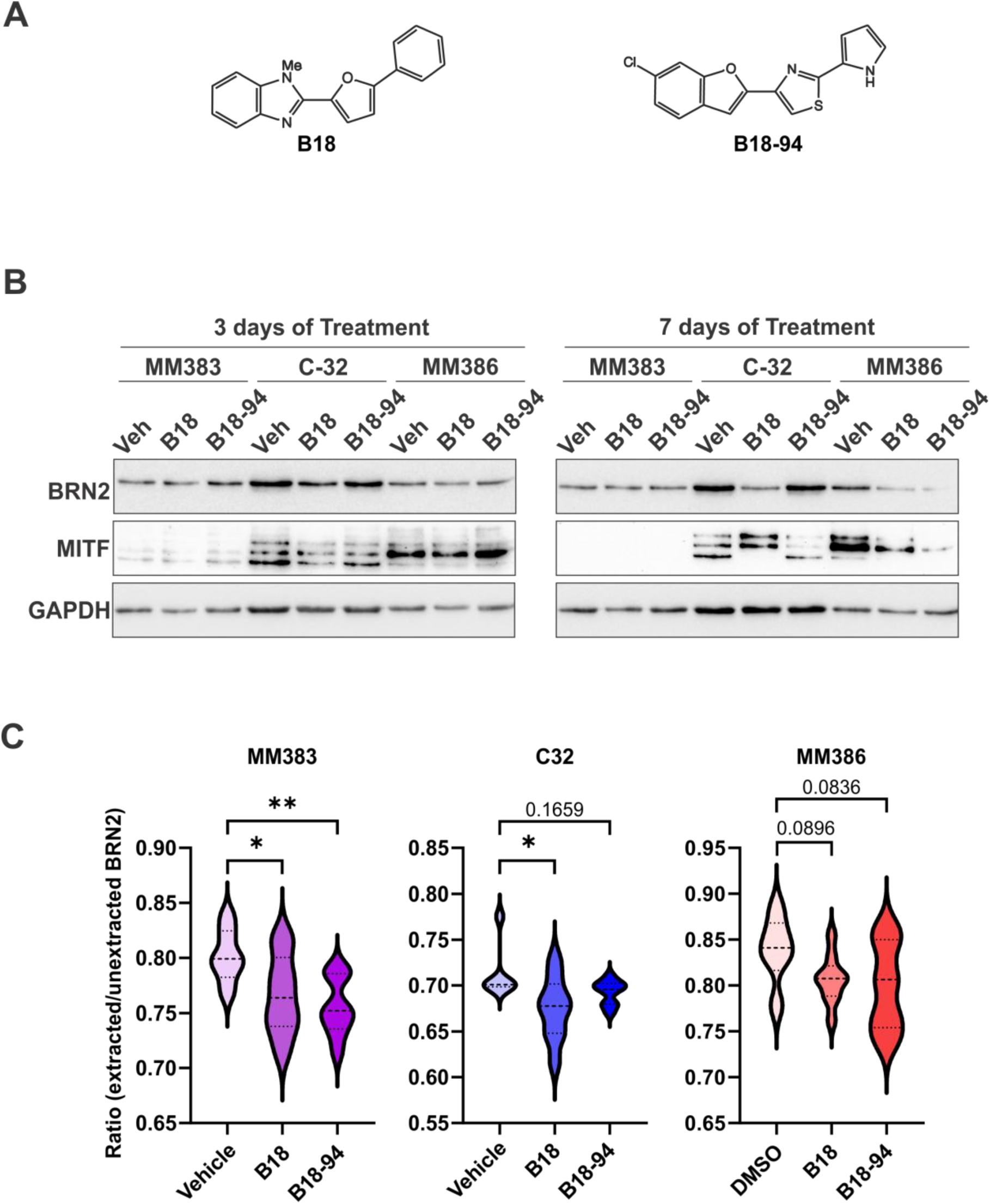
Small molecule inhibitors targeting the BRN2 transcription factor. (A) Structure of the BRN2 inhibitors. B18 and the B18-94 derivative as described in the Thaper *et al.* 2022 [49] preprint were examined in melanoma. (B) Western blot analysis following BRN2i treatment. Anoikis resistant cell lines (MM383, C-32 and MM386) were treated with 20 µM B18, 20 µM B18-94 or an equivalent volume of vehicle (DMSO) for 3 or 7 days in adherent culture. Protein was extracted and expression of BRN2, MITF and GAPDH as a loading control examined by western blotting. (C) Quantification of BRN2 binding to chromatin. Anoikis resistant cell lines were plated into 96 well glass bottom plates and treated with 20 µM B18, 20 µM B18-94 or vehicle. After 96 h, half of the wells for each treatment were pre-extracted, and then all wells fixed and permeabilized. Cells were primarily probed for BRN2 followed by an AlexaFluor-488 conjugated secondary antibody. Nuclei were stained with hoescht 33342 and cells imaged on the InCell Analyzer 6500 high content microscopy system. The ratio between pre-extracted and unextracted cells demonstrated reduced BRN2 chromatin binding following BRN2 inhibitor treatment. * = p <0.05, ** = p < 0.01, unpaired t-test with Welch’s correction for n = 3 independent biological replicates.

To confirm that the BRN2 inhibitors block the binding of BRN2 to DNA in melanoma cells, immunofluorescence was performed after treatment for 96 h **(Figure 2C, Supplementary Figure S3)**. The ratio between extracted and unextracted samples suggested a statistically significant reduction in BRN2 chromatin binding following treatment with B18 in both the MM383 (p = 0.0409) and C-32 (p = 0.0427) cell lines, and after treatment with B18-94 in MM383 cells (p = 0.0046) compared to vehicle. A trending reduction was observed after treatment with B18-94 in C-32 cells (p = 0.1659), and with both B18 (p = 0.0896) and B18-94 (p = 0.0836) in MM386 cells. These data indicated that the BRN2 inhibitors were able to reduce the binding of BRN2 to DNA in melanoma cell lines *in vitro*.

### BRN2 inhibition reduces cell survival in suspension culture

To examine the ability of the BRN2 inhibitors to reverse anoikis resistance in melanoma, a viability assay was performed by FACS following PI and calcein violet-AM staining **(Figure 3A)**. Treatment of anoikis resistant cell lines with 20 µM B18, or either 5 or 20 µM of B18-94 in suspension culture conditions revealed a significant reduction in percentage viability after 7 days. Treatment with B18 reduced the viability of MM383 cells by an average of 65.5 % (p = <0.0001), C-32 cells by 38.5 % (p = 0.0110) and MM386 cells by 25.5 % (p = 0.0028). In comparison, treatment with B18-94 resulted in a dose-dependent decrease in cell viability in the MM383 (42 % reduction at 20 µM, p = 0.0020) and C-32 cell lines (20 % reduction at 5 µM, p = 0.0301; 68 % reduction at 20 µM, p = 0.0133), but not in the MM386 cell line.

**Figure 3.**
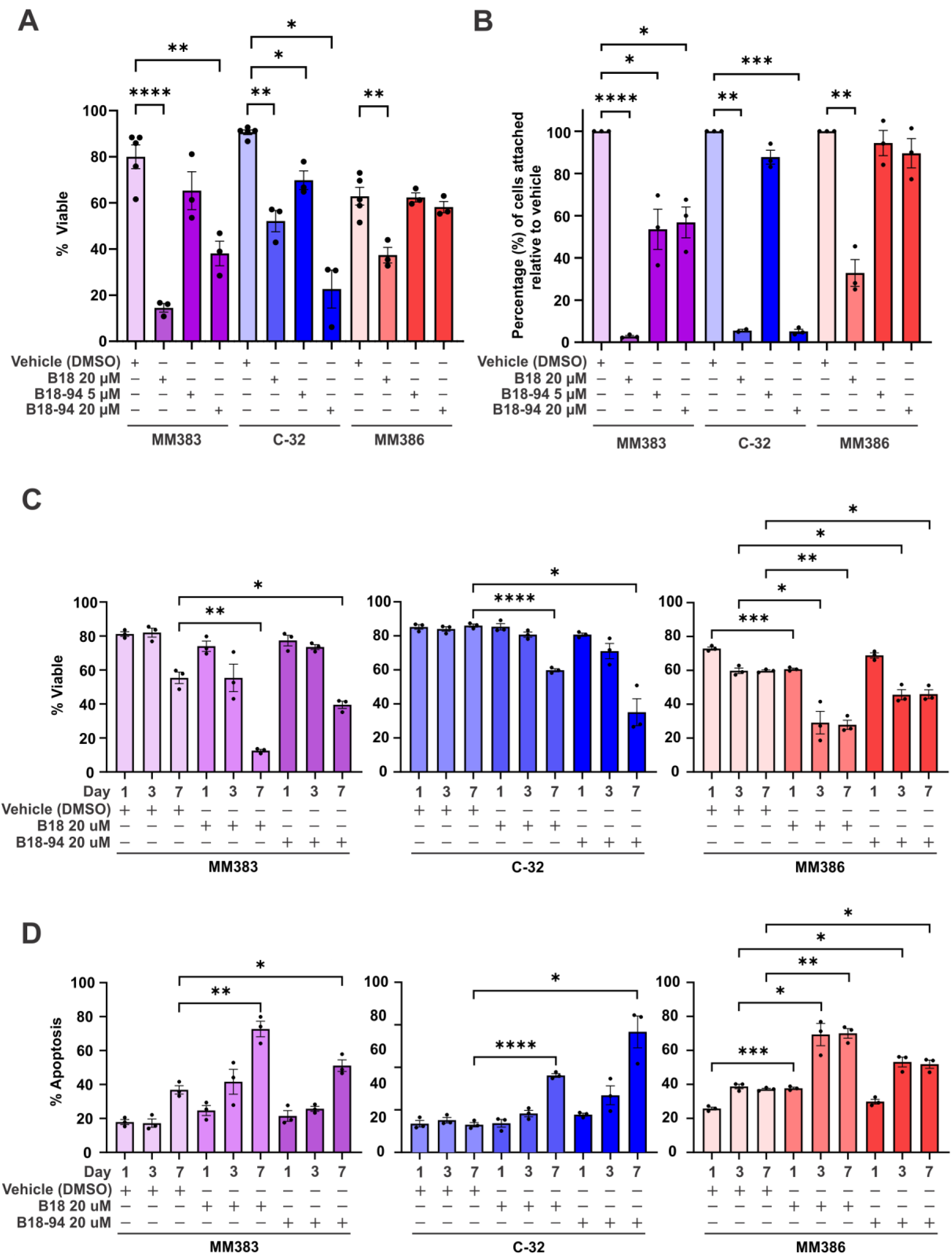
Inhibition of BRN2 reduces melanoma cell survival in suspension. MM383, C-32 and MM386 cell lines were plated into suspension culture conditions for 7 days with B18 (20 µM), B18-94 (5 or 20 µM), or vehicle (DMSO) added on days 0 and 4. (A) Cells were stained with 3 µM propidium iodide (PI) and 3 µM calcein violet-AM, and percentage (%) viability assessed by flow cytometry. (B) After treatment in suspension, cells were additionally transferred into adherent culture conditions for 72 h, and the proportion of cells able to reattach and grow quantitated by SRB assay. (C, D) Cells treated for 1, 3, or 7 days were stained with FITC-Annexin V and PI to observe (C) viable cells or (D) progression through the apoptotic cell death pathway. This revealed that BRN2 inhibition induces apoptosis in suspension conditions. Statistics presented for negative control versus drug treated for matching time points only. Values indicate mean ± standard error of the mean (SEM), n = 3 independent biological replicates, ((B) n = 2 – 3). * = p <0.05, ** = p < 0.01, *** = p <0.001, **** = p < 0.0001, unpaired t-test with Welch’s correction.

Following the 7 days of treatment in suspension culture, cells and drug-containing media were transferred into standard adherent culture. This confirmed that the BRN2 inhibitors significantly reduce the ability of cells to reattach and proliferate **(Figure 3B)**. After treatment with B18, the ability of MM383 cells to adhere was reduced by more than 97 % relative to the control (p = <0.0001). A 94 % reduction was observed in C-32 cells (p = 0.0040), with a 67 % reduction in MM386 cells (p = 0.0089). Treatment with B18-94 resulted in a significant reduction in cell attachment in the MM383 (46 % at 5 µM, p = 0.0399; 43 % at 20 µM, p = 0.0275) and C-32 cell lines only (94.88 % reduction at 20 µM, p = 0.0001), with no change observed in the MM386 cell line.

To further validate that the BRN2 inhibitors were inducing apoptosis in suspension culture conditions, cells were stained with Annexin-V and PI over a time course of treatment to measure changes in viability and track the passage of cells through either apoptosis or necrosis **(Figure 3C/D, Supplementary Figure S4)**. A significant reduction in cell viability and increase in apoptosis was confirmed following treatment with both B18 and B18-94 in all three cell lines. Compared to vehicle at 7 days of treatment, viability was reduced in MM383 cells by 42.9 % with B18 (p = 0.0043), and 15.9 % with B18-94 (p = 0.230), while apoptosis increased by 35.9 % (p = 0.0057) and 14.3 % (p = 0.0326), respectively. The C-32 cell line demonstrated an average 26.2 % reduction in viability with B18 (p = < 0.0001) and 50.9 % reduction with B18-94 (p = 0.0220), coinciding with an increase in apoptosis of 23.2 % (p = <0.0001) and 43.7 % (p = 0.0262). Comparatively, viability of MM386 cells reduced by 31.9 % with B18 (p = 0.0056) and 13.8 % with B18-94 (p = 0.0292) and apoptosis increased by 32.9 % (p = 0.0060) and 14.7 % (p = 0.0209). A negligible increase in necrosis was demonstrated at later time points and was statistically significant in the C-32 cell line only **(Supplementary Figure S4)**. These data highlight that the BRN2 inhibitors predominantly induce apoptosis, and suggests that these inhibitors function in blocking anoikis resistance.

### BRN2 drives mitochondrial dysfunction to prevent cell death by anoikis

To expand the understanding of the signaling pathways driving anoikis resistance downstream of BRN2, the BRN2 inhibitors were used as probes for analysis of the total proteome by Liquid Chromatography-Mass Spectrometry (LC-MS) **(Figure 4A, Supplementary File S5)**. As shown in **Figure 4B** and **Supplementary File S6**, IPA identified a range of signaling pathways consistently dysregulated between the three cell lines and both BRN2 inhibitor compounds. Notably, BRN2 inhibitor treatment demonstrated reduced mitochondrial dysfunction (average z-score = -4.58), PTEN signaling (-3.76) and PPAR signaling (-3.16), and increased oxidative phosphorylation (+6.59), mitochondrial translation (+7.26), and electron transport and ATP synthesis (+7.68) **(Figure 4C).** These pathway alterations suggested that BRN2 may drive anoikis resistance by dysregulating mitochondrial metabolism within the cell, dysregulating oxidative stress and reactive oxygen species (ROS) production, and subsequently the activity of apoptotic proteins. BRN2 inhibition was predicted to activate the MAPK and NF-κB signaling pathways by IPA, resulting in PPARɣ downregulation and restoring mitochondrial function, thereby allowing intrinsic cell death pathways to proceed **(Supplementary File S7)**.

**Figure 4.**
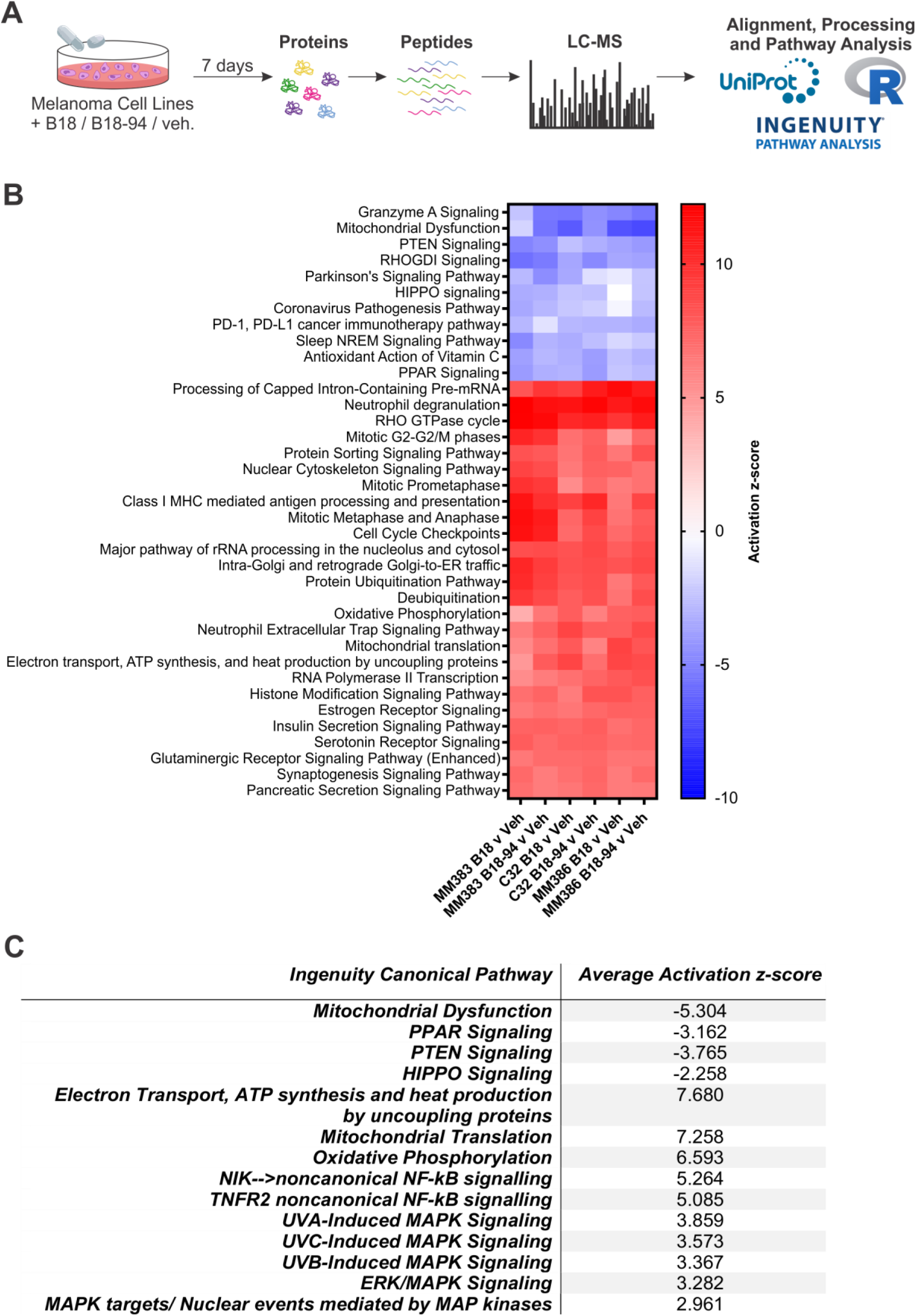
Proteomics analysis following BRN2 inhibition elucidates pathways of BRN2-mediated resistance to anoikis. (A) Workflow for proteomics analysis after BRN2i treatment. MM383, C-32 and MM386 cell lines were grown for 7 days in adherent culture with 20 µM B18, B18-94 or vehicle added on days 0 and 4. Cells were harvested and total proteome analyzed by Liquid Chromatography-Mass Spectrometry (LC-MS). Annotated proteomics data were analyzed for enrichment of canonical signaling pathways using Ingenuity Pathway Analysis (IPA) software after alignment and processing. (B) IPA analysis of proteomics data. Heatmap reveals variations in pathway enrichment for comparisons between treated and untreated samples for each of the cell lines. Activation z-score demonstrates pathway up- or down-regulation following hierarchical clustering. (C) Differentially expressed pathway signatures related to mitochondrial function. Activation z-scores within each pathway for the three cell lines and both B18 and B18-94 versus vehicle were averaged.

To validate the changes to mitochondrial activity observed following treatment with the BRN2 inhibitors, a MitoTracker Red FM assay was performed. Quantification of mitochondrial activity by reading MitoTracker Red FM fluorescence relative to hoescht 33342 staining revealed that treatment with B18 resulted in an approximately 30 % increase in mitochondrial activity after 3 days in all three cell lines **(Figure 5A)**. Following 7 days of treatment, the C-32 and MM386 cell lines demonstrated a 41 % (p = 0.0281) and 40 % (p = 0.0007) increase in mitochondrial activity respectively, while the effect of B18 in the MM383 cell line appeared unchanged between 3 and 7 days of treatment (22 % at 3 days; p = 0.0989). In comparison, treatment of all three cell lines with B18-94 had a minimal effect on mitochondrial activity at 3 days. After exposure to B18-94 for 7 days, the MM383 cell line demonstrated an increase in mitochondrial activity of 28 % (p = 0.0664), with a 20 % increase observed for both C-32 (p = 0.0194) and MM386 (p = 0.0809).

**Figure 5.**
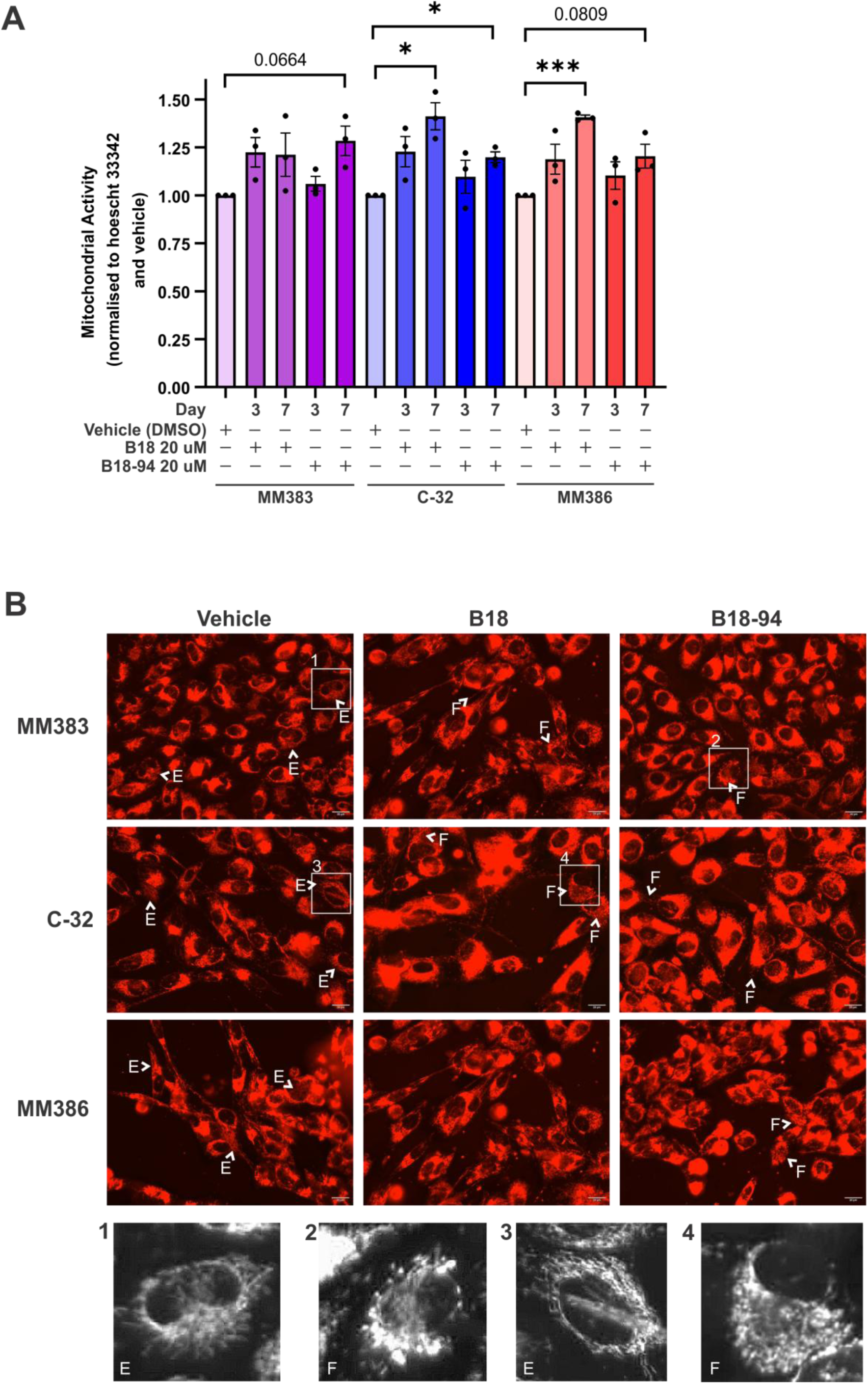
BRN2 regulates mitochondrial dysfunction to drive anoikis resistance. To validate the mitochondrial phenotype identified by proteomics, MM383, C-32 and MM386 cells were grown with B18 (20 µM), B18-94 (20 µM), or vehicle (DMSO) for 3 or 7 days and then mitochondria stained with MitoTracker Red FM and nuclei stained with hoescht 33342. (A) Mitochondrial activity was quantitated on the Biotek Synergy Neo2 fluorescent plate reader and normalized to hoechst 33342 fluorescence and vehicle after 3 and 7 days of treatment. This revealed an increase in mitochondrial function with BRN2 inhibition. Values indicate mean ± standard error of the mean (SEM), n = 3 independent biological replicates. * = p <0.05, *** = p <0.001, unpaired t-test with Welch’s correction. (B) Representative images after 7 days of treatment were taken on the Evos FL microscope at 40× magnification. White arrows indicate examples of increased mitochondrial fragmentation observed upon BRN2 inhibition, E = elongated mitochondrial morphology, F = fragmented mitochondrial staining indicative of increased ROS production and mitochondrial fission. Scale bars = 20 µm. Numbered boxes indicate cropped greyscale images.

In addition to the increase in mitochondrial activity quantitated by the fluorescent plate reader, mitochondrial morphology was observed by fluorescent microscopy following 7 days of treatment **(Figure 5B)**. While mitochondria in the vehicle treated cells appeared to be largely elongated, these images revealed treatment resulted in an increase in mitochondrial fragmentation as indicated by granular or punctate staining. Overall, the MitoTracker Red FM assay confirmed that inhibition of BRN2 resulted in an increase in mitochondrial activity in all three cell lines over time.

### BRN2 inhibition sensitizes cells to killing by vemurafenib

To examine whether small molecule inhibition of BRN2 was able to sensitize cells to BRAF-targeted therapy, the MM383, C-32 and MM386 cell lines were treated with the BRN2 inhibitors in combination with increasing concentrations of the BRAF-targeted inhibitor vemurafenib, in adherent or non-adherent culture conditions **(Figure 6)**.

**Figure 6.**
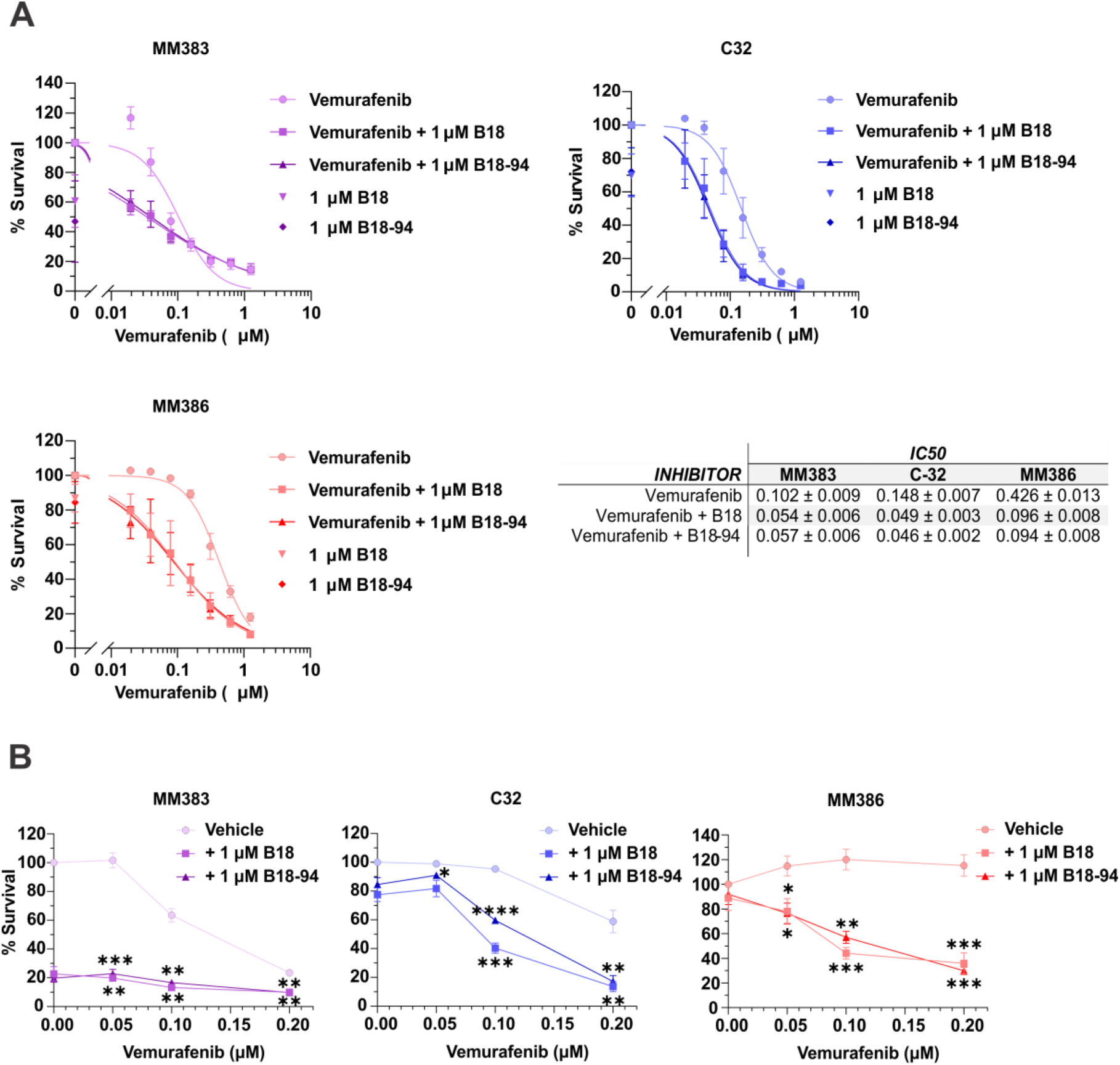
BRN2 inhibitors sensitize cells to killing by BRAF inhibition. To examine whether the BRN2 inhibitors were able to sensitize to BRAF inhibition, MM383, C-32 and MM386 cell lines were seeded into (A) adherent (5,000 cells per well) or (B) ultra-low attachment (ULA; 20,000 cells per well) conditions and treated with 1 µM B18 or B18-94 with or without increasing concentrations of vemurafenib. After 7 days, cell survival was quantitated by SRB assay. For ULA plates, cells were transferred to standard coated 96-well plates with an extra 100 µL media, incubated for 24 hours before SRB assay. (A) Percentage survival relative to vemurafenib alone was calculated in Microsoft Excel and IC_50_ calculated with a variable slope non-linear fit model for inhibitor vs. normalized response in GraphPad Prism. Values indicate IC_50_ ± standard error of the mean (SEM), n = 6 independent biological replicates. (B) Values indicate percentage survival relative to vemurafenib alone, mean ± SEM, n = 3-4 independent biological replicates. * = p <0.05, ** = p <0.01, *** = p <0.001, **** = p <0.0001, unpaired t-test with Welch’s correction.

In adherent culture, the combination of vemurafenib with 1 µM of either B18 or B18-94 was able to reduce the IC_50_ value relative to vemurafenib alone in all three cell lines **(Figure 6A)**. In MM383 cells, the combination treatment halved the IC_50_ from 0.102 ± 0.009 µM with vemurafenib alone to 0.054 ± 0.006 µM in combination with B18 and 0.057 ± 0.006 µM in combination with B18-94. In C-32 cells, the IC_50_ was reduced from 0.148 ± 0.007 µM to 0.049 ± 0.003 µM with B18 and 0.046 ± 0.002 µM with B18-94. While the IC_50_ was reduced from 0.426 ± 0.013 µM to 0.096 ± 0.008 µM with B18 and 0.094 ± 0.008 µM with B18-94 in the MM386 cell line.

In ultra-low attachment conditions, the combination treatments significantly reduced the percentage of cells surviving relative to vemurafenib treatment alone **(Figure 6B)**. In MM383 cells, treatment with 1 µM of either B18 or B18-94 reduced survival by 78 %. Due to the sensitivity of these cells to BRN2 inhibition alone, the effect of the combination with BRAF inhibition was subtle, with a further reduction in survival to 9 % with 0.2 µM vemurafenib (p = 0.0071 for B18, p = 0.0050 for B18-94). In the C-32 cell line, BRN2 inhibition alone resulted in a 16 % (B18) and 23 % (B18-94) reduction in percentage survival. The combination with 0.1 µM vemurafenib reduced survival by 60 % (B18, p < 0.0001) and 40 % (B18-94, p = 0.0003), while the combination with 0.2 µM reduced survival by 87 % (B18, p = 0.0056) and 83 % (B18-94, p = 0.0068). In MM386 cells, BRN2 inhibition alone reduced survival by 12 %. The combination with 0.05 µM vemurafenib reduced survival by 24 % with both B18 (p = 0.0337) and B18-94 (p = 0.0164), BRN2 inhibition with 0.1 µM vemurafenib reduced survival by 56 % (B18, p = 0.0007) and 43 % (B18-94, p = 0.0015), and the combination with 0.2 µM vemurafenib further reduced survival by 65 % (B18, p = 0.0006) and 71 % (B18-94, p = 0.0010).

Overall, the BRN2 inhibitors sensitized human melanoma cell lines to killing by targeted inhibition of BRAF using vemurafenib in both adherent and non-adherent culture conditions.

## 4.0 Discussion

Accumulating evidence suggests that the BRN2 *(POU3F2)* transcription factor is an important contributor to the progression of malignant melanoma to metastasis. High levels of BRN2 expression have been demonstrated to induce an invasive phenotype, characterized by slow cycling, highly motile cells with stem-cell like properties [34, 35]. Our laboratory identified that BRN2 has a role in promoting resistance to cell death by anoikis in melanoma [40], where engineered over-expression of BRN2 resulted in a significant increase in viability of cells cultured in non-adherent conditions. However, the mechanisms driving this phenomenon downstream of BRN2 remained to be elucidated. Given the reciprocal relationship between MITF and BRN2 expression controlling phenotypic plasticity [37, 41–43], our study confirmed that both MITF downregulation and BRN2 function in anoikis resistance.

Despite its role in disease progression, targeting BRN2 with small molecule inhibitors has not been investigated in melanoma. Thaper *et al.* described the development of BRN2 small molecule inhibitors that interact with the POU-S and POU-H DNA binding domains and inhibit transcription factor binding and expression of invasive signature genes in neuro-endocrine prostate cancer cells [49]. By applying these small molecule BRN2 inhibitors, we demonstrated that BRN2 is a druggable target for the killing of anoikis resistant melanoma cells in non-adherent culture conditions, and sensitizes cells to killing by vemurafenib. Further development of the BRN2 inhibitors may improve the efficacy of their application to specifically target disseminated cells in circulation, preventing the seeding of metastases. Specifically, Kobi *et al.* described that in melanoma, BRN2 preferentially binds DNA containing MORE (More Palindromic Octamer Factor Recognition Element) motif sequences with 0, 1 or 2 base insertions (ATRNATATRCAW, ATRNATnATRCAW or ATRNATnnATRCAW) [57]. Conversely, the BRN2 inhibitors designed by Thaper *et al.* were created to target BRN2 binding to alternate MORE motif sequences (GTATGCAAATAAG) [49]. Subsequently, the efficacy and specificity of the BRN2 inhibitors may be improved by targeting BRN2 binding to the specific MORE motifs.

Moreover, we utilized the BRN2 inhibitors as probes to better understand the BRN2 signaling cascade in anoikis resistance. Through analysis of the total proteome following BRN2 inhibitor treatment, we identified that BRN2 dysregulates mitochondrial metabolism, oxidative stress and reactive oxygen species (ROS) production, and subsequently the activity of apoptotic proteins through the MAPK and NF-κB signaling pathways and PPARγ expression. PPARγ is known to control both intrinsic and extrinsic cell death pathways by decreasing Bcl-2 expression and increasing pro-apoptotic Bax by regulating ROS generation [58]. In melanoma, PPARγ has been demonstrated to control migration downstream of FAK by acting as a regulator of STAT3 activity via miR-125b [58]. STAT3, a known regulator of anoikis resistance, is shown to be activated downstream of BRN2 in suspension culture [40]. Subsequently, the regulation of PPARγ by BRN2 would be expected to have implications on mitochondrial function and metabolism, and the regulation of apoptosis [59].

A reduction in ROS production by NADPH during oxidative phosphorylation, and an increase in ROS scavenging, have been shown to perpetuate circulating tumor cell (CTC) survival [60, 61], and aligns with the increase in ROS inferred by the proteomics data following BRN2 inhibition. Furthermore, downregulation of glucose metabolism and upregulation of fatty acid oxidation (FAO) have been shown to reduce excessive ROS through production of antioxidants nicotinamide adenine dinucleotide and flavin adenine dinucleotide in melanoma [62]. In our study, the predicted increase in oxidative phosphorylation observed following BRN2 inhibition coincided with an increase in activation of both glucose metabolism and FAO pathways **(Supplementary File S6)**. The upregulation of proteins for fatty acid transport and β-oxidation, in particular CROT and CRAT, have been demonstrated to contribute to anoikis resistance and correlate with reduced overall survival of melanoma patients, while the use of FAO inhibitors such as thioridazine and ranolazine reduce colony formation of melanoma cells [62]. Other studies have shown that melanoma CTCs regulate FAO via PGC1α [63, 64], reducing ROS by activating anti-ferroptosis programs [65] and managing oxidative stress by upregulating autophagy [66]. However, whether an increase or decrease in ROS drives anoikis resistance appears to be highly context dependent, differing between cancer types [60, 67, 68]. Contradicting research suggest that upregulation of ROS such as hydrogen peroxide drives anoikis resistance by upregulating caveolin-1 (Cav-1), resulting in the downstream activation of AKT signaling and cell survival [20, 69].

Using MitoTracker Red FM, we confirmed that treatment with the BRN2 inhibitors reduced mitochondrial dysfunction, observed through an increase in mitochondrial activity and mitochondrial fragmentation or fission [70]. Mitochondrial dysfunction is an important adaptation acquired by cancer cells that allows a robust response to persistent stress conditions such as hypoxia, metabolic and proliferative stress, driving survival and plasticity [71, 72], as well as resistance to apoptosis and cancer therapies [73]. An increase in mitochondrial fragmentation or fission has been reported to assist in the clearing of dysfunctional mitochondria and is an important early step allowing the intrinsic apoptotic cascade to proceed [74–76]. Fission can be triggered by ROS, and results in mitochondrial outer membrane permeabilization, cytochrome c release and caspase activation [77].

Mitochondrial dysfunction in cancer cells has been proposed as a therapeutic target, where the restoration of mitochondrial function leads to the release of pro-apoptotic factors and caspase activation, eventuating in programmed cell death [78]. Existing inhibitors that reduce mitochondrial dysfunction have been shown to increase glucose oxidation and induce ROS, initiating cell death in a range of cancer models [79]. Conversely, treatment of melanoma cells with the BRAF inhibitor, vemurafenib, results in mitochondrial hyperfusion by blocking fission and enhancing fusion components [80]. This process is suggested to be a resistance mechanism to vemurafenib treatment, while the targeting of hyperfusion was demonstrated to enhance its cytotoxicity. Importantly, BRN2 expression has been associated with resistance to BRAF inhibition in a range of studies [40, 81–83]. The sensitivity of melanoma cells to MAPK inhibitor therapy was further linked to metabolic signatures driven by PPARγ [84], while alterations in the level of ROS and oxidative damage have been demonstrated in MAPKi resistant melanomas [85]. Here, we demonstrated that combining BRN2 inhibitors with the BRAF-targeted inhibitor vemurafenib was able to sensitize cells to killing by vemurafenib in both adherent and non-adherent culture conditions. This suggests that the restoration of mitochondrial function by the BRN2 inhibitors may prevent the mitochondrial hyperfusion induced by vemurafenib and has the potential to reverse resistance to MAPK-targeted therapy.

In summary, we confirm a role for BRN2 in anoikis resistance and demonstrated that targeting BRN2 with small molecule inhibitors induced apoptosis in melanoma cells under non-adherent conditions through the reversal of mitochondrial dysfunction. Small molecule inhibition of BRN2, in combination with existing MAPK-targeted therapies, may allow the targeting of metastatic cells in circulation, and prevent the seeding of metastatic lesions, improving the prognosis for melanoma patients.

## 5.0 Acknowledgements

We wish to acknowledge the contribution of the QIMR Berghofer sample processing, sequencing and proteomics facilities to this project.

## 6.0 Author contributions

HMN and GMB wrote the manuscript. HMN, XH, MNA and GMB performed the experiments. HMN, MNA, KAT, JLS and GMB analyzed the data. HMN, JLS, AGS, PVB, CMW and GMB were responsible for conceptualization. All authors read and reviewed the manuscript.

## Supplementary Figures

**Figure S1.**
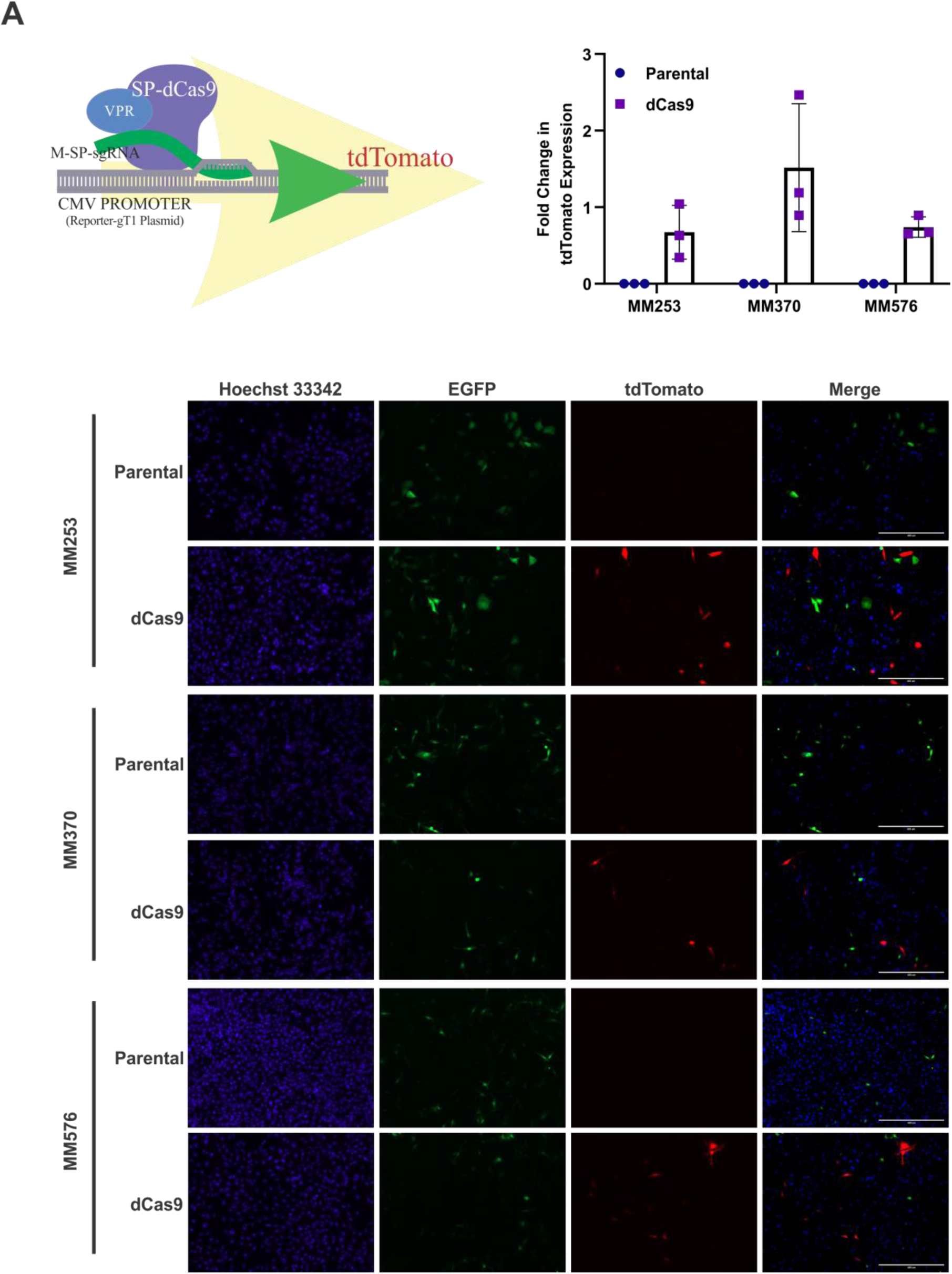
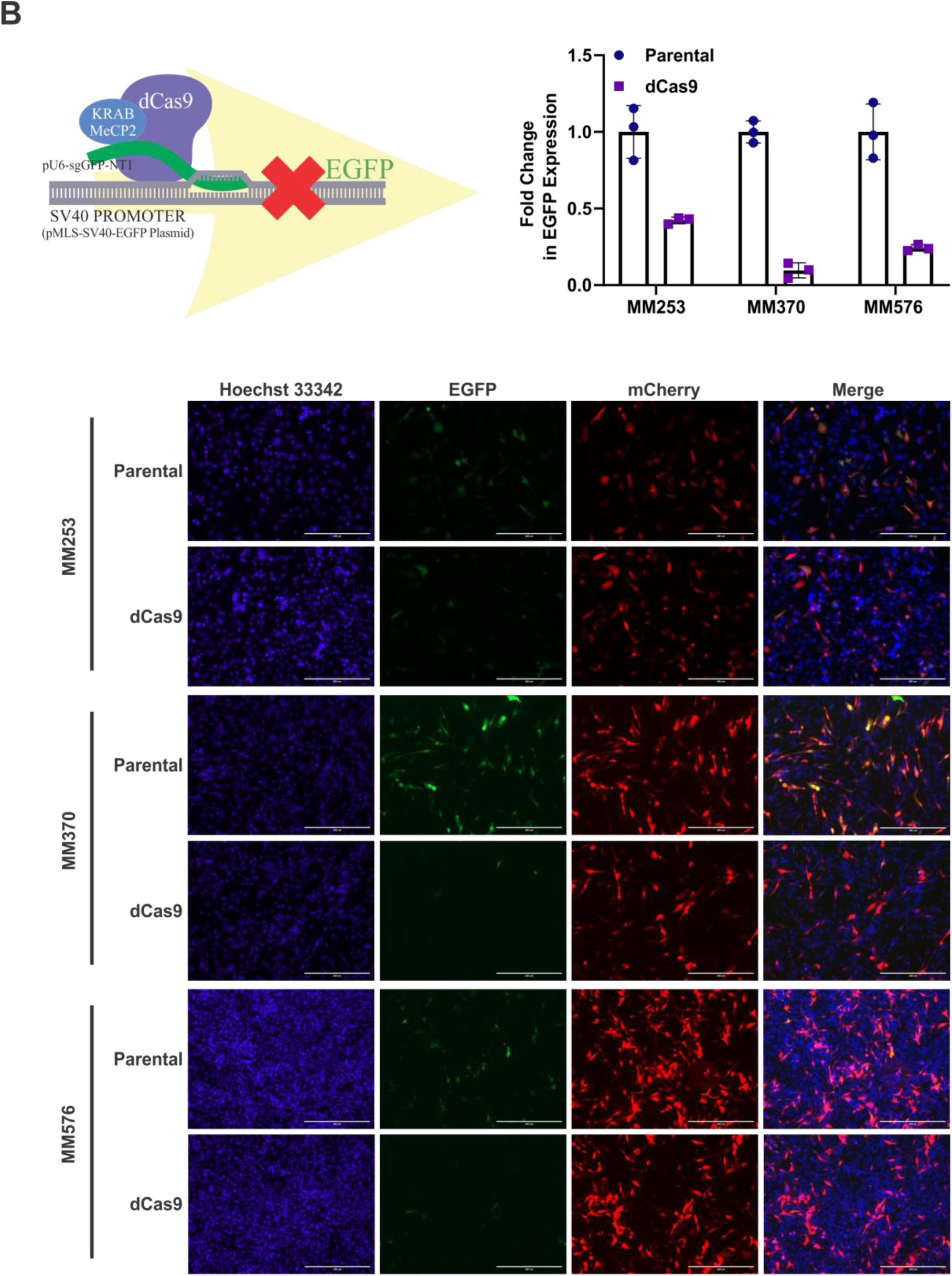
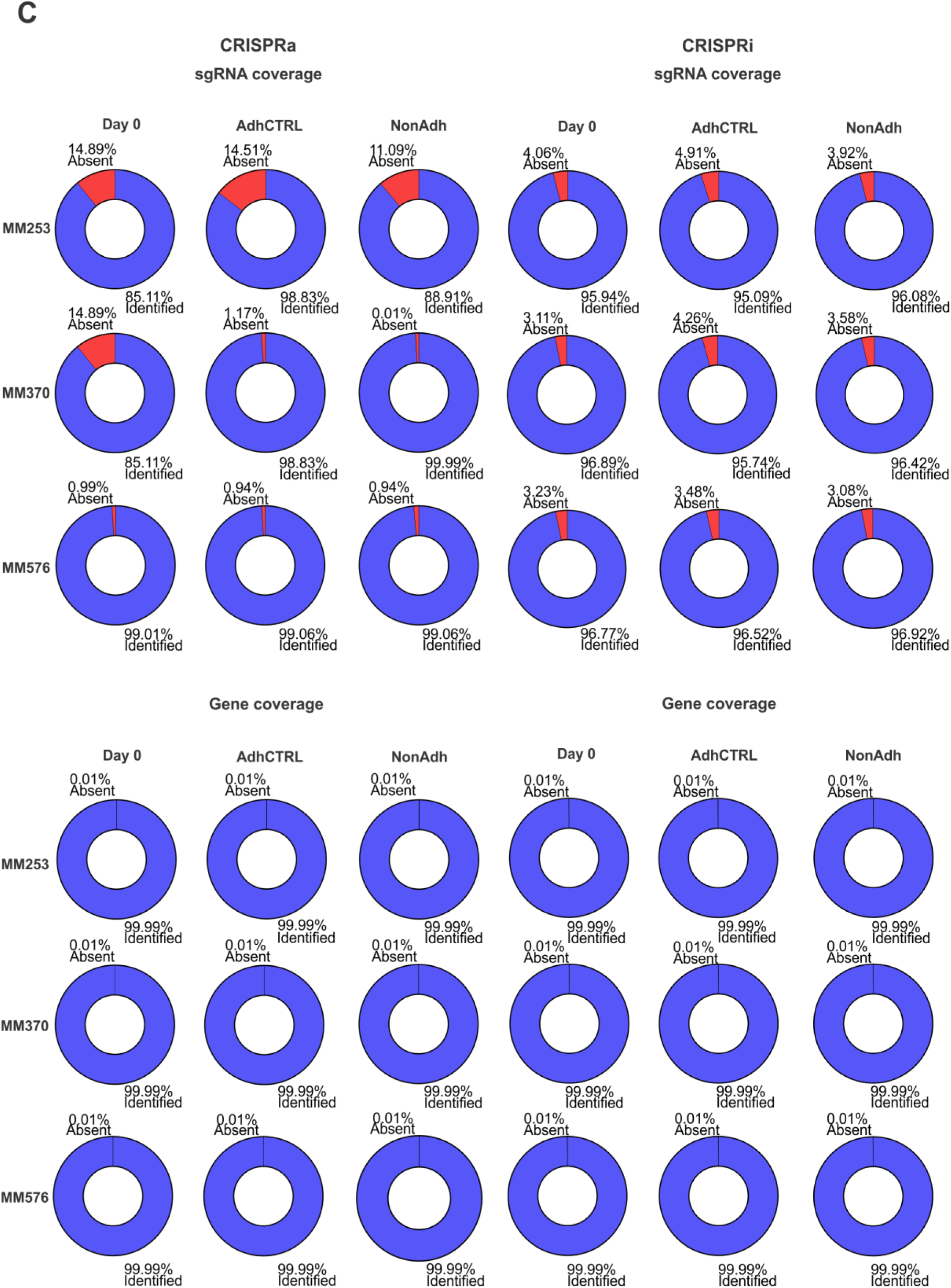
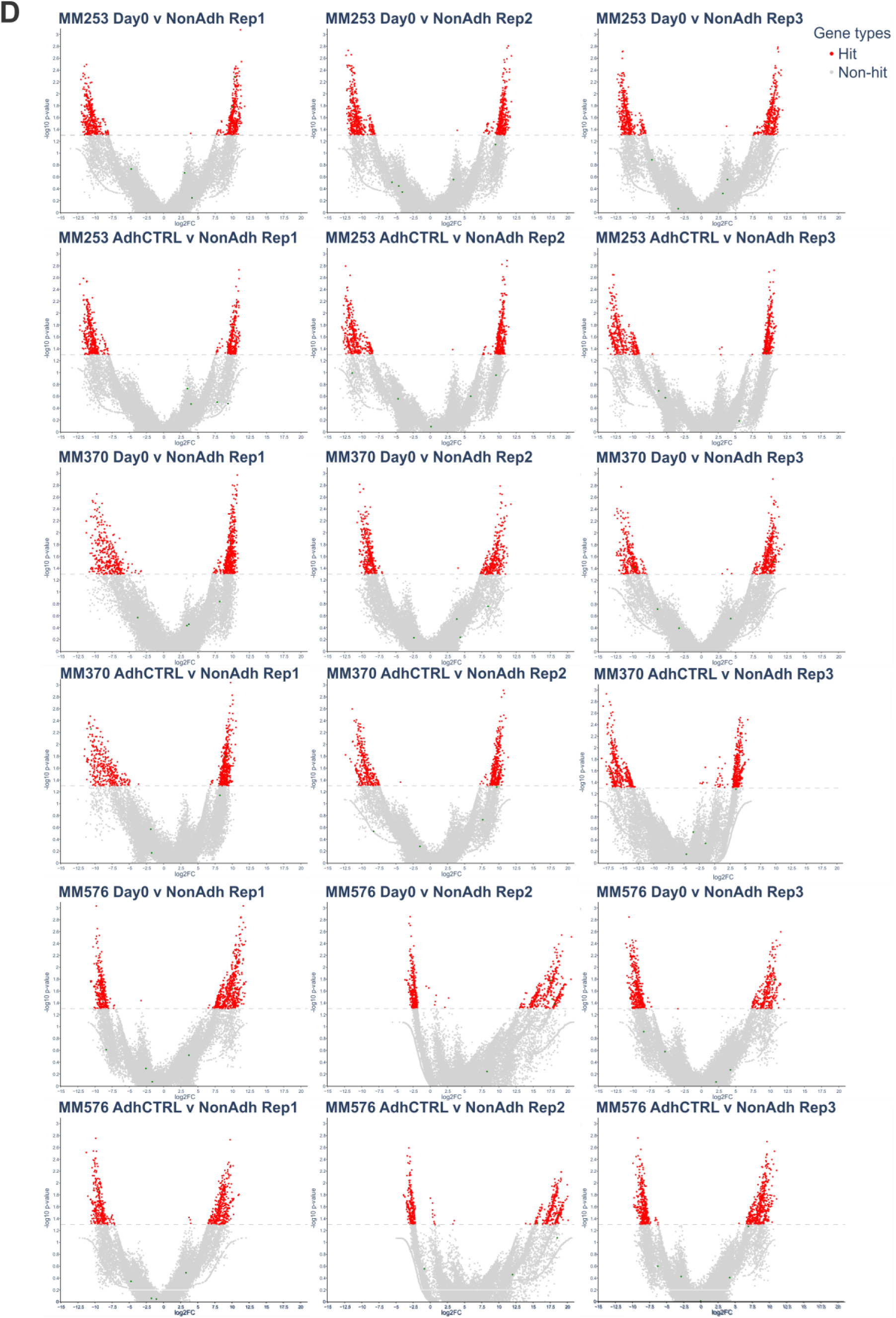
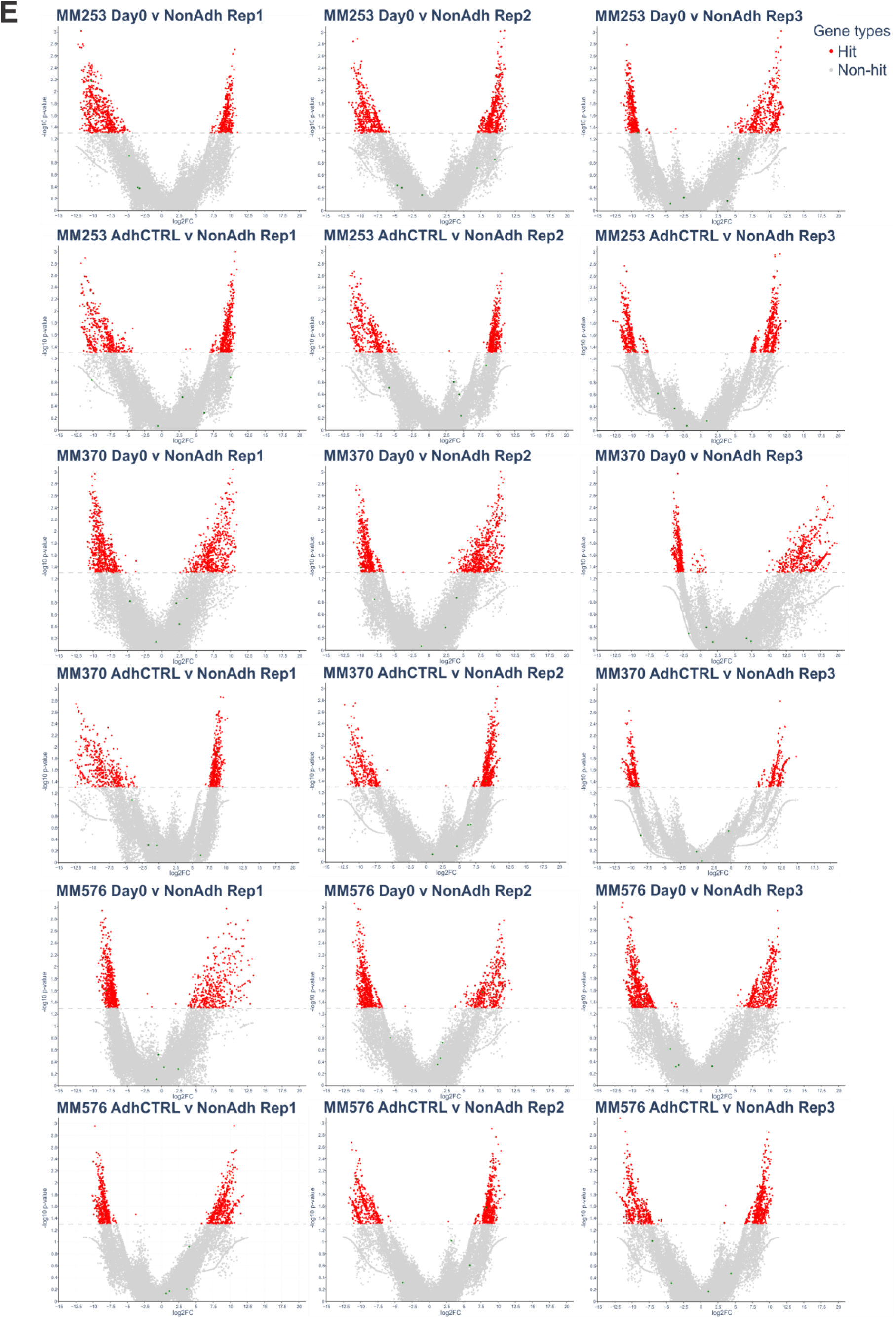
Additional CRISPR screening data. (A/B) To validate dCas9 activity, reporter assays were performed. (A) For CRISPRa, parental and SP-dCas9-VPR expressing cells were co-transfected with the reporter-gT1, M-SP-sgRNA and pLenti6-EGFP plasmids. (B) For CRISPRi, parental and lenti_dCas9-KRAB-MeCP2 expressing cells were co-transfected with the pMLS-SV40-EGFP and pU6-sgGFP-NT1 plasmids. Cells were cultured for 48 h after transfection, stained with 30 µg/mL hoechst 33342, and cells imaged on the EVOS FL Auto Microscope. Quantification of fluorescent cells was performed in QuPath-0.2.3 software. Fold-change in reporter expression (dCas9 activity) was calculated accounting for the ratio, transfection efficiency, and probability of achieving co-transfection of sgRNA and reporter plasmids. Parental cell line = blue circles; dCas9 cell line = purple squares, n = 3 random fields of view, averaged. Images present one representative field of view per sample. Scale bar = 400 µm, images taken with 10× objective lens. (C) Pie charts show sgRNA and gene coverage for day 0, adherent control (AdhCTRL), and non-adherent (NonAdh) samples for the MM253, MM370 and MM576 cell lines in both the CRISPRa and CRISPRi screens. Coverage was calculated from counts generated by the Horlbeck *et al.* ScreenProcessing Python pipeline. Counts were then merged across the three replicates, and percentages calculated for each sample using python. (D/E) Volcano plots showing gene enrichment between the Day 0 and NonAdh, or AdhCTRL and NonAdh samples for individual replicates in the (D) CRISPRa and (E) CRISPRi screens as log_2_ fold-change plotted against –log_10_ Mann-Whitney p-value. Dashed line = 0.05 p-value, grey points = non-hit genes, red points = significant genes.

**Figure S2.**
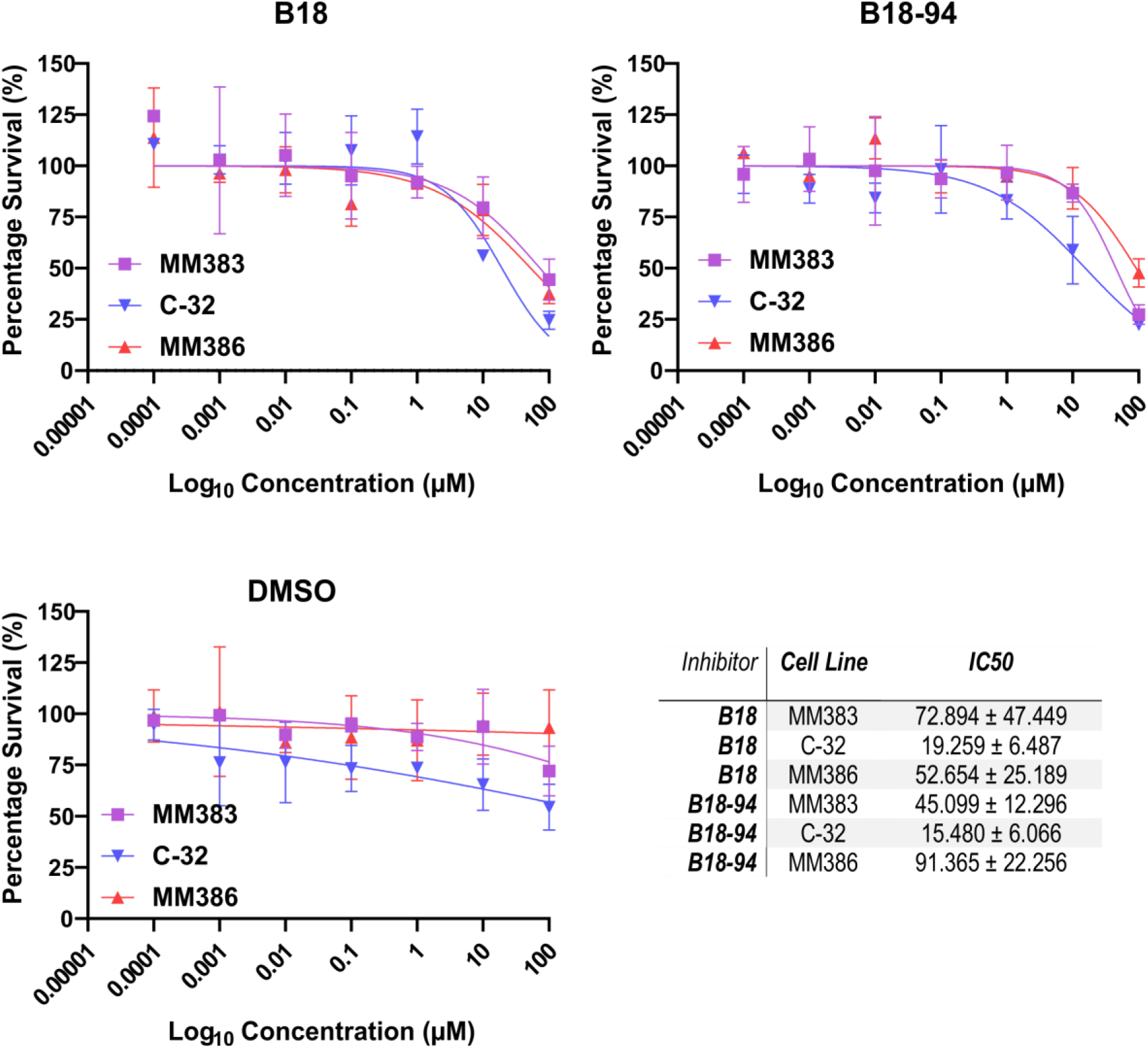
IC50 for BRN2 inhibitors in adherent culture. MM383 (purple squares), C-32 (blue inverted triangles) and MM386 (orange triangles) cell lines treated with 10-fold serial dilutions of B18, B18-94 or equivalent volumes of DMSO from 100 µM to 0.0001 µM in triplicate. Cell viability was quantitated by SRB assay after 7 days. Percentage survival relative to negative control was calculated in Microsoft Excel and IC_50_ calculated with a variable slope non-linear fit model for inhibitor vs. normalized response in GraphPad Prism. Values indicate IC_50_ ± standard error of the mean (SEM), n = 2 independent biological replicates.

**Figure S3.**
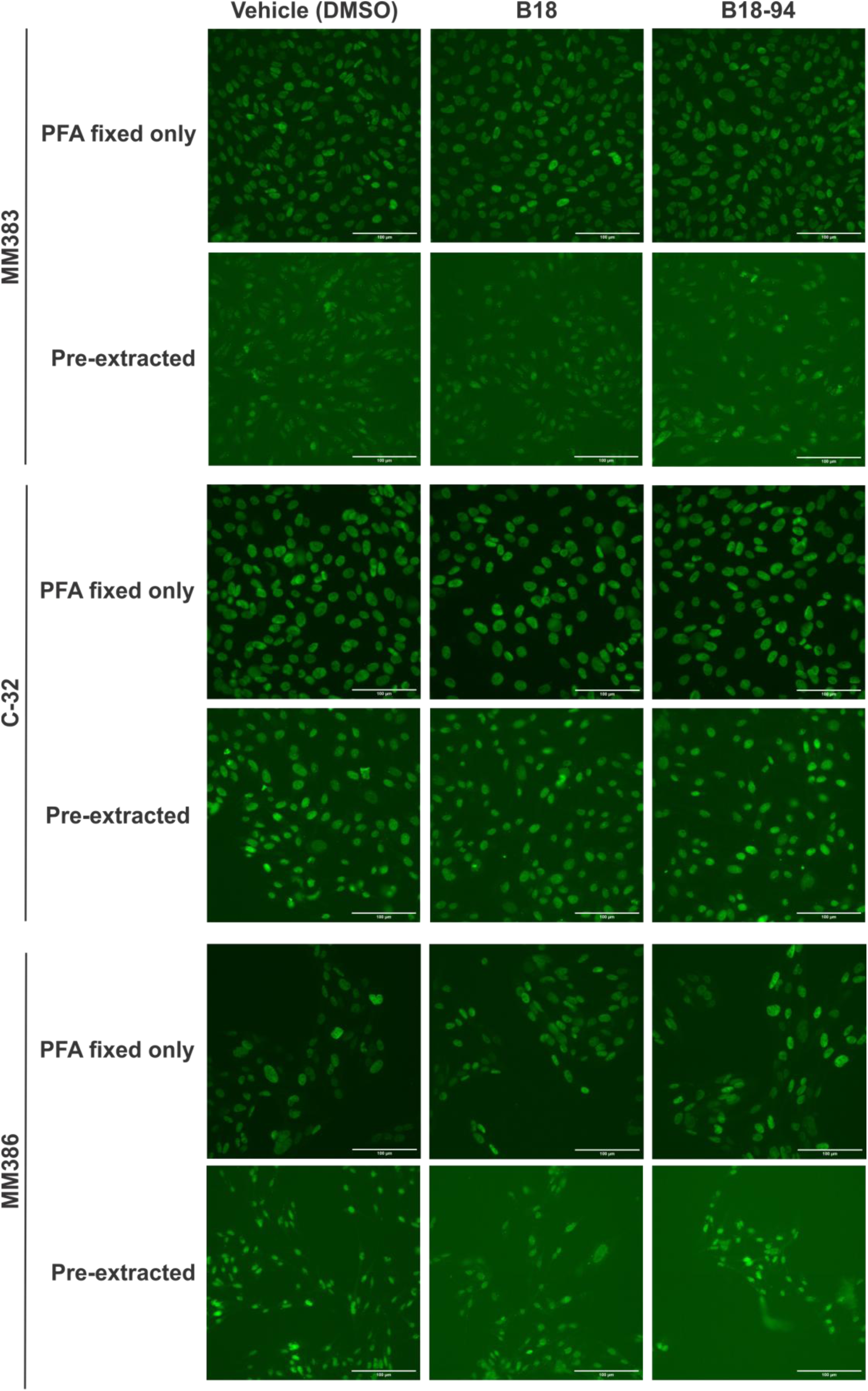
Representative images of BRN2 inhibitor immunofluorescence. Representative microscopy images for quantification of BRN2 binding to chromatin for each cell line and treatment type (40× magnification). Anoikis resistant cell lines were plated into 96 well glass bottom plates and treated with 20 µM B18, 20 µM B18-94 or vehicle. After 96 h, half of the wells for each treatment were pre-extracted, and then the plate fixed and permeabilized. Cells were primarily probed for BRN2 followed by an AlexaFluor-488 conjugated secondary antibody. Nuclei were stained with hoescht 33342 and cells imaged on the InCell Analyzer 6500 high content microscopy system.

**Figure S4.**
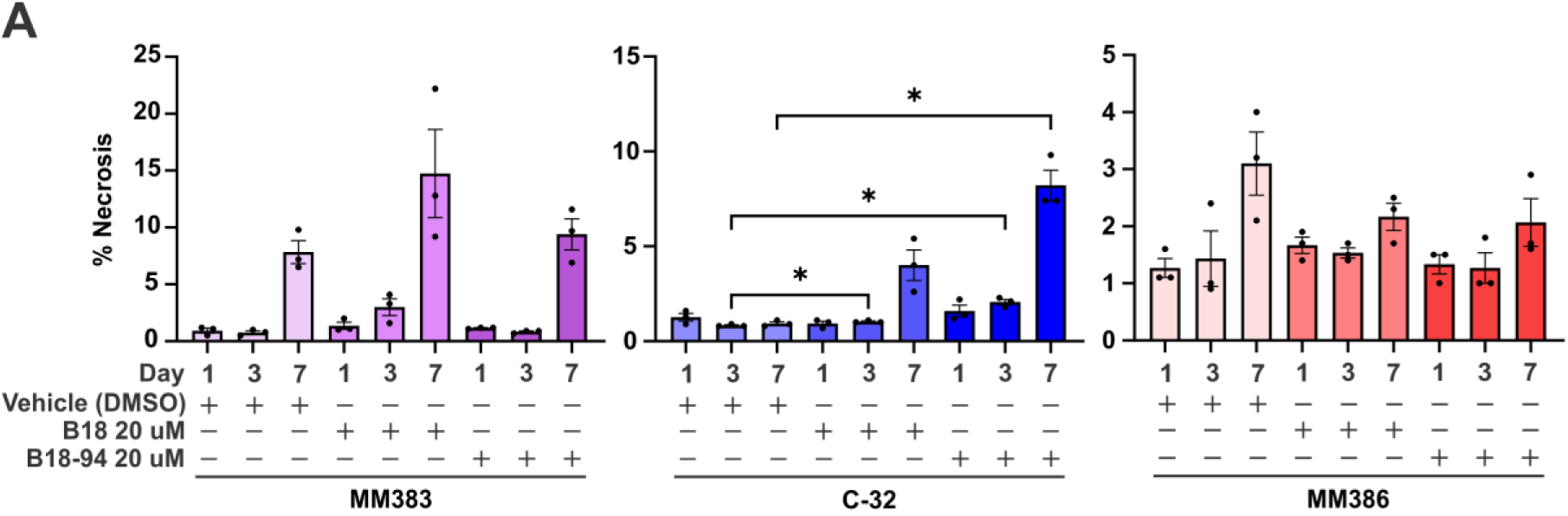
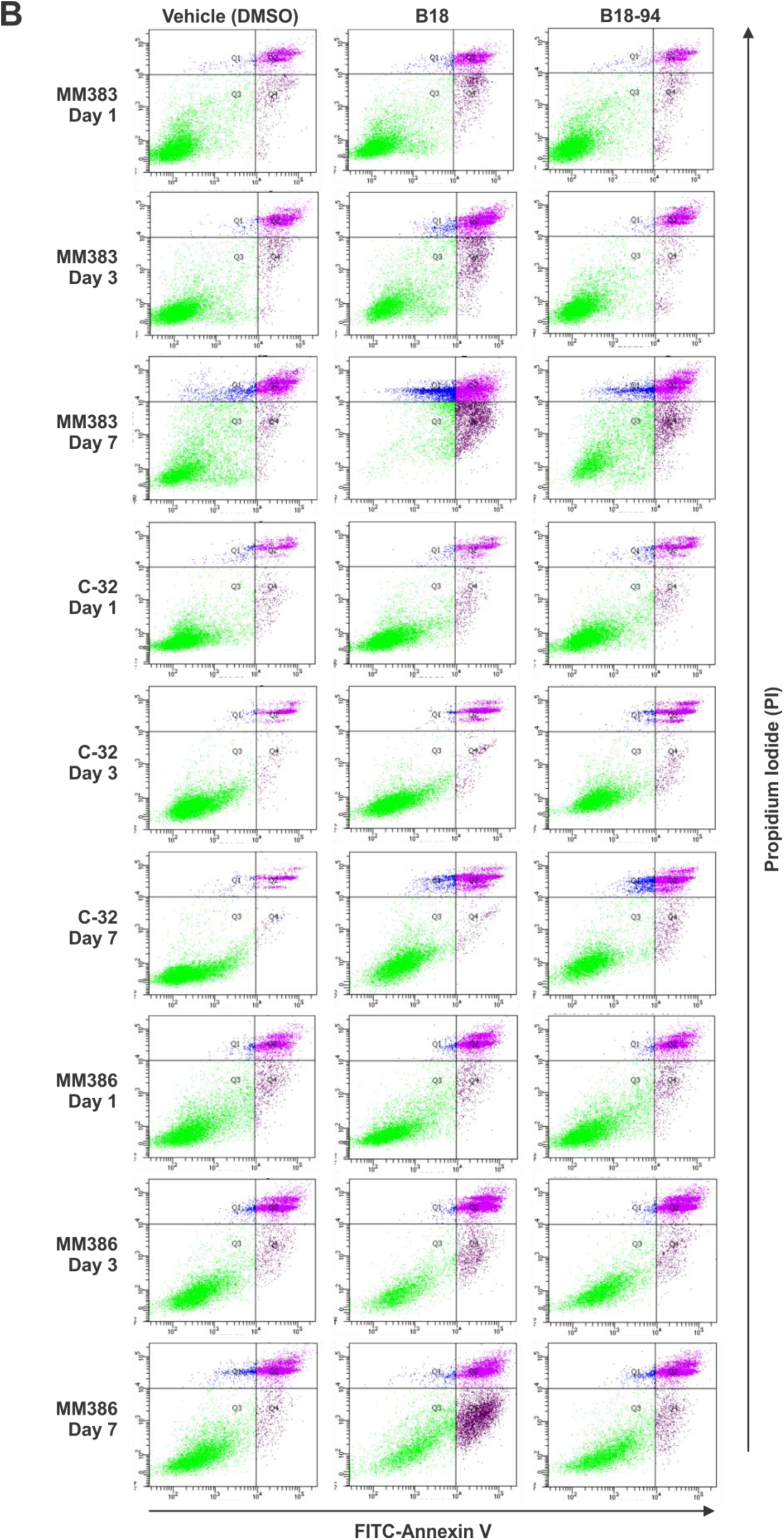
BRN2 inhibitor flow cytometry. MM383, C-32 and MM386 cell lines were plated into suspension culture conditions for 1, 3 or 7 days with B18 (20 µM), B18-94 (20 µM), or vehicle (DMSO) added on days 0 and 4. Cells were stained with FITC-Annexin V and propidium iodide (PI) as described in Figure 3 and (A) percentage (%) necrosis calculated. (B) Apoptosis in the graphs is the sum of the early and late apoptosis quadrants (Q2 and Q4). Q1 are necrotic cells, and Q3 are live cells. FACS plots demonstrate one representative replicate per timepoint for each cell line. (A) Statistics presented for negative control versus drug treated for matching time points only. Values indicate mean ± standard error of the mean (SEM), n = 3 independent biological replicates. * = p <0.05, ** = p < 0.01, *** = p <0.001, **** = p < 0.0001, unpaired t-test with Welch’s correction.

**Supplementary File S1**

Synthetic procedures and characterization, including ^1^H NMR spectra plus ORTEP diagrams and unit cells for B18 and B18-94.

**Supplementary File S2**

Excel file of CRISPRa screening results presented as a gene table of comparisons between the Day 0, adherent control (AdhCTRL) and non-adherent (NonAdh) samples for each replicate performed, as well as the average of the replicates (best_avg) for the MM253, MM370 and MM576 cell lines. Significantly altered genes in each comparison based on Mann-Whitney p-value < 0.05.

**Supplementary File S3**

Excel file of CRISPRi screening results presented as a gene table of comparisons between the Day 0, adherent control (AdhCTRL) and non-adherent (NonAdh) samples for each replicate performed, as well as the average of the replicates (best_avg) for the MM253, MM370 and MM576 cell lines. Significantly altered genes in each comparison based on Mann-Whitney p-value < 0.05.

**Supplementary File S4**

Excel file for Ingenuity Pathway Analysis (IPA) results as of 28 Apr, 2024 for significantly altered genes compiled from both the CRISPRa and CRISPRi screens.

**Supplementary File S5**

Excel file of all protein enrichment from comparisons between BRN2 inhibitor and vehicle treated samples for the MM383, C-32 and MM386 cell lines as determined by proteomics analysis.

**Supplementary File S6**

Excel file for comparisons of activation z-score between samples for canonical pathways enriched in proteomics following BRN2 inhibitor treatment.

**Supplementary File S7**

Excel file for differentially expressed proteins for mitochondrial canonical pathways consistently altered in cell lines treated with B18 or B18-94 as determined by IPA.

## References

1. Liu, C., et al., Global, regional, and national burden of cutaneous malignant melanoma from 1990 to 2021 and prediction to 2045. Front Oncol, 2024. 14: p. 1512942.

2. Brenner, M. and V.J. Hearing, The protective role of melanin against UV damage in human skin. Photochem Photobiol, 2008. 84(3): p. 539–49.

3. Sample, A. and Y.Y. He, Mechanisms and prevention of UV-induced melanoma. Photodermatol Photoimmunol Photomed, 2018. 34(1): p. 13–24.

4. Davis, L.E., S.C. Shalin, and A.J. Tackett, Current state of melanoma diagnosis and treatment. Cancer Biol Ther, 2019. 20(11): p. 1366–1379.

5. von Schuckmann, L.A., et al., Risk of Melanoma Recurrence After Diagnosis of a High-Risk Primary Tumor. JAMA Dermatol, 2019. 155(6): p. 688–693.

6. Bollag, G., et al., Vemurafenib: the first drug approved for BRAF-mutant cancer. Nat Rev Drug Discov, 2012. 11(11): p. 873–86.

7. Wright, C.J. and P.L. McCormack, Trametinib: first global approval. Drugs, 2013. 73(11): p. 1245–54.

8. Menzies, A.M. and G.V. Long, Dabrafenib and trametinib, alone and in combination for BRAF-mutant metastatic melanoma. Clin Cancer Res, 2014. 20(8): p. 2035–43.

9. FDA, New Medical Devicescollab. P t, 2015. 40(12): p. 792–806.

10. Lipson, E.J. and C.G. Drake, Ipilimumab: an anti-CTLA-4 antibody for metastatic melanoma. Clin Cancer Res, 2011. 17(22): p. 6958–62.

11. Robert, C., et al., Five-Year Outcomes With Nivolumab in Patients With Wild-Type BRAF Advanced Melanoma. J Clin Oncol, 2020. 38(33): p. 3937–3946.

12. Barone, A., et al., FDA Approval Summary: Pembrolizumab for the Treatment of Patients with Unresectable or Metastatic Melanoma. Clin Cancer Res, 2017. 23(19): p. 5661–5665.

13. Albrecht, L.J., et al., The Latest Option: Nivolumab and Relatlimab in Advanced Melanoma. Curr Oncol Rep, 2023. 25(6): p. 647–657.

14. Wolchok, J.D., et al., Final, 10-Year Outcomes with Nivolumab plus Ipilimumab in Advanced Melanoma. N Engl J Med, 2025. 392(1): p. 11–22.

15. Haugh, A.M., A.K.S. Salama, and D.B. Johnson, Advanced Melanoma: Resistance Mechanisms to Current Therapies. Hematol Oncol Clin North Am, 2021. 35(1): p. 111–128.

16. Frisch, S. and H. Francis, Disruption of epithelial cell-matrix interactions induces apoptosis. Journal of Cell Biology, 1994. 124(4): p. 619–626.

17. Frisch, S.M. and R.A. Screaton, Anoikis mechanisms. Curr Opin Cell Biol, 2001. 13(5): p. 555–62.

18. Meredith, J.E., Jr., B. Fazeli, and M.A. Schwartz, The extracellular matrix as a cell survival factor. Mol Biol Cell, 1993. 4(9): p. 953–61.

19. Mei, J., et al., Anoikis in cell fate, physiopathology, and therapeutic interventions. MedComm (2020), 2024. 5(10): p. e718.

20. Paoli, P., E. Giannoni, and P. Chiarugi, Anoikis molecular pathways and its role in cancer progression. Biochim Biophys Acta, 2013. 1833(12): p. 3481–3498.

21. Frisch, S.M., M. Schaller, and B. Cieply, Mechanisms that link the oncogenic epithelial-mesenchymal transition to suppression of anoikis. J Cell Sci, 2013. 126(Pt 1): p. 21–9.

22. Vandamme, N. and G. Berx, From neural crest cells to melanocytes: cellular plasticity during development and beyond. Cell Mol Life Sci, 2019. 76(10): p. 1919–1934.

23. Pedri, D., et al., Epithelial-to-mesenchymal-like transition events in melanoma. The FEBS Journal, 2022. 289(5): p. 1352–1368.

24. Neuendorf, H.M., J.L. Simmons, and G.M. Boyle, Therapeutic targeting of anoikis resistance in cutaneous melanoma metastasis. Frontiers in Cell and Developmental Biology, 2023. 11.

25. Boisvert-Adamo, K. and A.E. Aplin, B-RAF and PI-3 kinase signaling protect melanoma cells from anoikis. Oncogene, 2006. 25(35): p. 4848–4856.

26. Boisvert-Adamo, K. and A.E. Aplin, Mutant B-RAF mediates resistance to anoikis via Bad and Bim. Oncogene, 2008. 27(23): p. 3301–12.

27. Boisvert-Adamo, K., et al., Mcl-1 is required for melanoma cell resistance to anoikis. Mol Cancer Res, 2009. 7(4): p. 549–56.

28. Fofaria, N.M. and S.K. Srivastava, Critical role of STAT3 in melanoma metastasis through anoikis resistance. Oncotarget, 2014. 5(16): p. 7051–64.

29. Peppicelli, S., et al., Anoikis Resistance as a Further Trait of Acidic-Adapted Melanoma Cells. J Oncol, 2019. 2019: p. 8340926.

30. Shao, Y. and A.E. Aplin, Akt3-mediated resistance to apoptosis in B-RAF-targeted melanoma cells. Cancer Res, 2010. 70(16): p. 6670–81.

31. Cook, A.L., et al., Human Melanoblasts in Culture: Expression of BRN2 and Synergistic Regulation by Fibroblast Growth Factor-2, Stem Cell Factor, and Endothelin-3. Journal of Investigative Dermatology, 2003. 121(5): p. 1150–1159.

32. Eisen, T., et al., The POU domain transcription factor Brn-2: elevated expression in malignant melanoma and regulation of melanocyte-specific gene expression. Oncogene, 1995. 11(10): p. 2157–64.

33. Thomson, J.A., et al., The brn-2 gene regulates the melanocytic phenotype and tumorigenic potential of human melanoma cells. Oncogene, 1995. 11(4): p. 691–700.

34. Goodall, J., et al., The Brn-2 transcription factor links activated BRAF to melanoma proliferation. Mol Cell Biol, 2004. 24(7): p. 2923–31.

35. Zeng, H., et al., Bi-allelic Loss of CDKN2A Initiates Melanoma Invasion via BRN2 Activation. Cancer Cell, 2018. 34(1): p. 56–68.e9.

36. Perego, M., et al., A slow-cycling subpopulation of melanoma cells with highly invasive properties. Oncogene, 2018. 37(3): p. 302–312.

37. Fane, M.E., et al., NFIB Mediates BRN2 Driven Melanoma Cell Migration and Invasion Through Regulation of EZH2 and MITF. EBioMedicine, 2017. 16: p. 63–75.

38. Thurber, A.E., et al., Inverse expression states of the BRN2 and MITF transcription factors in melanoma spheres and tumour xenografts regulate the NOTCH pathway. Oncogene, 2011. 30(27): p. 3036–48.

39. Pinner, S., et al., Intravital imaging reveals transient changes in pigment production and Brn2 expression during metastatic melanoma dissemination. Cancer Res, 2009. 69(20): p. 7969–77.

40. Pierce, C.J., et al., BRN2 expression increases anoikis resistance in melanoma. Oncogenesis, 2020. 9(7): p. 64.

41. Simmons, J.L., et al., MITF and BRN2 contribute to metastatic growth after dissemination of melanoma. Sci Rep, 2017. 7(1): p. 10909.

42. Goodall, J., et al., Brn-2 Represses Microphthalmia-Associated Transcription Factor Expression and Marks a Distinct Subpopulation of Microphthalmia-Associated Transcription Factor–Negative Melanoma Cells. Cancer Research, 2008. 68(19): p. 7788–7794.

43. Boyle, G.M., et al., Melanoma cell invasiveness is regulated by miR-211 suppression of the BRN2 transcription factor. Pigment Cell & Melanoma Research, 2011. 24(3): p. 525–537.

44. Stark, M. and N. Hayward, Genome-Wide Loss of Heterozygosity and Copy Number Analysis in Melanoma Using High-Density Single-Nucleotide Polymorphism Arrays. Cancer Research, 2007. 67(6): p. 2632–2642.

45. Pavey, S., et al., Microarray expression profiling in melanoma reveals a BRAF mutation signature. Oncogene, 2004. 23(23): p. 4060–7.

46. Gill, K.P. and M. Denham, Optimized Transgene Delivery Using Third-Generation Lentiviruses. Current Protocols in Molecular Biology, 2020. 133(1): p. e125.

47. Horlbeck, M.A., et al., Compact and highly active next-generation libraries for CRISPR-mediated gene repression and activation. Elife, 2016. 5.

48. Boyle, G.M., et al., Macrophage inhibitory cytokine-1 is overexpressed in malignant melanoma and is associated with tumorigenicity. J Invest Dermatol, 2009. 129(2): p. 383–91.

49. Thaper, D., et al., Discovery and characterization of a first-in-field transcription factor BRN2 inhibitor for the treatment of neuroendocrine prostate cancer. bioRxiv, 2022: p. 2022.05.04.490172.*preprint

50. Kildey, K., et al., Elevating CDCA3 levels in non-small cell lung cancer enhances sensitivity to platinum-based chemotherapy. Communications Biology, 2021. 4(1): p. 638.

51. Shah, E.T., et al., Inhibition of Aurora B kinase (AURKB) enhances the effectiveness of 5-fluorouracil chemotherapy against colorectal cancer cells. British Journal of Cancer, 2024. 130(7): p. 1196–1205.

52. Pino, L.K., et al., Acquiring and Analyzing Data Independent Acquisition Proteomics Experiments without Spectrum Libraries. Mol Cell Proteomics, 2020. 19(7): p. 1088–1103.

53. Demichev, V., et al., DIA-NN: neural networks and interference correction enable deep proteome coverage in high throughput. Nat Methods, 2020. 17(1): p. 41–44.

54. Jones, J., et al., Tidyproteomics: an open-source R package and data object for quantitative proteomics post analysis and visualization. BMC Bioinformatics, 2023. 24(1): p. 239.

55. Carreira, S., et al., Mitf regulation of Dia1 controls melanoma proliferation and invasiveness. Genes Dev, 2006. 20(24): p. 3426–39.

56. Hoek, K.S. and C.R. Goding, Cancer stem cells versus phenotype-switching in melanoma. Pigment Cell Melanoma Res, 2010. 23(6): p. 746–59.

57. Kobi, D., et al., Genome-wide analysis of POU3F2/BRN2 promoter occupancy in human melanoma cells reveals Kitl as a novel regulated target gene. Pigment Cell & Melanoma Research, 2010. 23(3): p. 404–418.

58. Pei, G., et al., FAK regulates E-cadherin expression via p-SrcY416/p-ERK1/2/p-Stat3Y705 and PPARγ/miR-125b/Stat3 signaling pathway in B16F10 melanoma cells. Oncotarget, 2017. 8(8): p. 13898–13908.

59. Tol, M.J., et al., A PPARγ-Bnip3 Axis Couples Adipose Mitochondrial Fusion-Fission Balance to Systemic Insulin Sensitivity. Diabetes, 2016. 65(9): p. 2591–605.

60. Adeshakin, F.O., et al., Targeting Oxidative Phosphorylation-Proteasome Activity in Extracellular Detached Cells Promotes Anoikis and Inhibits Metastasis. Life (Basel), 2021. 12(1).

61. Zhu, G., et al., Metastatic Melanoma Cells Rely on Sestrin2 to Acquire Anoikis Resistance via Detoxifying Intracellular ROS. J Invest Dermatol, 2020. 140(3): p. 666–675.e2.

62. Lasheras-Otero, I., et al., The Regulators of Peroxisomal Acyl-Carnitine Shuttle CROT and CRAT Promote Metastasis in Melanoma. J Invest Dermatol, 2023. 143(2): p. 305–316.e5.

63. Vazquez, F., et al., PGC1α expression defines a subset of human melanoma tumors with increased mitochondrial capacity and resistance to oxidative stress. Cancer Cell, 2013. 23(3): p. 287–301.

64. Luo, C., et al., A PGC1α-mediated transcriptional axis suppresses melanoma metastasis. Nature, 2016. 537(7620): p. 422–426.

65. Hong, X., et al., The Lipogenic Regulator SREBP2 Induces Transferrin in Circulating Melanoma Cells and Suppresses Ferroptosis. Cancer Discov, 2021. 11(3): p. 678–695.

66. Fung, C., et al., Induction of autophagy during extracellular matrix detachment promotes cell survival. Mol Biol Cell, 2008. 19(3): p. 797–806.

67. Giannoni, E., et al., Redox regulation of anoikis: reactive oxygen species as essential mediators of cell survival. Cell Death & Differentiation, 2008. 15(5): p. 867–878.

68. Du, S., et al., NADPH oxidase 4 regulates anoikis resistance of gastric cancer cells through the generation of reactive oxygen species and the induction of EGFR. Cell Death & Disease, 2018. 9(10): p. 948.

69. Halim, H. and P. Chanvorachote, Long-term hydrogen peroxide exposure potentiates anoikis resistance and anchorage-independent growth in lung carcinoma cells. Cell Biol Int, 2012. 36(11): p. 1055–66.

70. Neikirk, K., et al., MitoTracker: A useful tool in need of better alternatives. European Journal of Cell Biology, 2023. 102(4): p. 151371.

71. O’Malley, J., et al., Mitochondrial Stress Response and Cancer. Trends Cancer, 2020. 6(8): p. 688–701.

72. Fendt, S.M., C. Frezza, and A. Erez, Targeting Metabolic Plasticity and Flexibility Dynamics for Cancer Therapy. Cancer Discov, 2020. 10(12): p. 1797–1807.

73. Hsu, C.C., L.M. Tseng, and H.C. Lee, Role of mitochondrial dysfunction in cancer progression. Exp Biol Med (Maywood), 2016. 241(12): p. 1281–95.

74. Chen, W., H. Zhao, and Y. Li, Mitochondrial dynamics in health and disease: mechanisms and potential targets. Signal Transduction and Targeted Therapy, 2023. 8(1): p. 333.

75. Suen, D.F., K.L. Norris, and R.J. Youle, Mitochondrial dynamics and apoptosis. Genes Dev, 2008. 22(12): p. 1577–90.

76. Montessuit, S., et al., Membrane Remodeling Induced by the Dynamin-Related Protein Drp1 Stimulates Bax Oligomerization. Cell, 2010. 142(6): p. 889–901.

77. Ježek, J., K.F. Cooper, and R. Strich, The Impact of Mitochondrial Fission-Stimulated ROS Production on Pro-Apoptotic Chemotherapy. Biology (Basel), 2021. 10(1).

78. Chen, G., et al., Preferential killing of cancer cells with mitochondrial dysfunction by natural compounds. Mitochondrion, 2010. 10(6): p. 614–25.

79. Bhat, T.A., et al., Restoration of mitochondria function as a target for cancer therapy. Drug Discov Today, 2015. 20(5): p. 635–43.

80. Ferraz, L.S., et al., Targeting mitochondria in melanoma: Interplay between MAPK signaling pathway and mitochondrial dynamics. Biochemical Pharmacology, 2020. 178: p. 114104.

81. Herbert, K., et al., BRN2 suppresses apoptosis, reprograms DNA damage repair, and is associated with a high somatic mutation burden in melanoma. Genes Dev, 2019. 33(5-6): p. 310–332.

82. Smith, M.P., et al., A PAX3/BRN2 rheostat controls the dynamics of BRAF mediated MITF regulation in MITF(high) /AXL(low) melanoma. Pigment Cell Melanoma Res, 2019. 32(2): p. 280–291.

83. Zhang, Y., et al., Bidirectional interconversion between mutually exclusive tumorigenic and drug-tolerant melanoma cell phenotypes. bioRxiv, 2023: p. 2020.08.26.269126.

84. Zhang, G., et al., Targeting mitochondrial biogenesis to overcome drug resistance to MAPK inhibitors. J Clin Invest, 2016. 126(5): p. 1834–56.

85. Pizzimenti, S., et al., Oxidative Stress-Related Mechanisms in Melanoma and in the Acquired Resistance to Targeted Therapies. Antioxidants (Basel), 2021. 10(12).

