## Supplementary File S1 for "Inhibition of BRN2 in Melanoma Reverses Anoikis Resistance and Sensitizes Cells to Killing by Vemurafenib"

### Synthetic Procedures and Characterization for B18 and B18-94

#### General experimental

Glassware were oven dried before use with anhydrous solvents and reagents. Tetrahydrofuran was freshly distilled from elemental sodium / benzophenone under an argon atmosphere. Dimethyl sulfoxide (DMSO) was freshly distilled to dryness over calcium hydride (4% w/v) under reduced pressure and stored over activated molecular sieves (4Å) under an argon atmosphere.

All commercially available reagents and chemicals were purified where possible, except for Xantphos which was used as received. Reaction, extraction and chromatography solvents, e.g. petroleum spirits (boiling point range 40 – 60 °C), ethyl acetate, ethanol (EtOH), dimethoxy ethane (DME), and dichloromethane (DCM) were freshly distilled before use.

$^{1}$H NMR spectra were recorded using a Bruker Avance 400 MHz spectrometer at 25 °C. Chemical shifts (δ) are reported in parts per million (ppm) and referenced internally according to residual solvent peaks: $CDCl_3$ ($^{1}$H δ: 7.26 ppm – $CHCl_3$), DMSO-$d_6$ ($^{1}$H δ: 2.50 ppm – DMSO) and acetone-$d_6$ ($^{1}$H δ: 2.17 ppm - acetone). The following abbreviations are used to report multiplicities: s = singlet, d = doublet, t = triplet, q = quartet, m = multiplet.

For thin-layer chromatography (TLC), Merck aluminum plates coated with 60 F254 silica gel were employed and were visualized under UV light ($\lambda_{max}$ = 254 nm). Merck LC60A 40-30 silica gel was used for column chromatography.

#### Experimental procedures

##### 1-Methyl-2-(5-phenylfuran-2-yl)-1*H*-benzo[*d*]imidazole (B18)

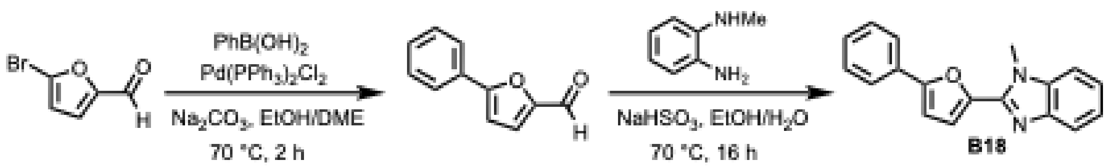


5-Bromo-2-furaldehyde (0.55 g, 3.14 mmol), phenylboronic acid (0.49 g, 4.01 mmol), and 2M $Na_2CO_3$ (5 mL) were added to a solution of ethanol (8 mL) and dimethoxyethane (4 mL). The mixture was then purged with argon for 20 min. Subsequently, $PdCl_2(PPh_3)_2$ (0.20 g 0.28 mmol) was added to the reaction mixture, which was again purged with argon for 10 min at room temperature. The reaction mixture was heated to 70 °C under an argon atmosphere. After 2 h the reaction mixture was cooled to room temperature and diluted with water (25 mL) followed by extraction with ethyl acetate (3 x 25 mL). The organic layers were washed with water (25 mL), dried over anhydrous $Na_2SO_4$, and filtered. The filtrate was concentrated under reduced pressure, and the crude subjected to silica gel column chromatography (20% ethyl acetate in petroleum spirit) to afford 5-phenylfuran-2-carbaldehyde as a pale-yellow oil (0.53 g, 98%). Spectroscopic data were in accordance with the literature[1]. $^1$H-NMR (400 MHz, $CDCl_3$): δ (ppm) 9.66 (s, 1H), 7.85-7.80 (m, 2H), 7.48-7.39 (m, 3H), 7.33-7.31 (d, 1H), 6.86-6.83 (d, 1H).

Following a modified procedure of Jiang *et al*[2]: 5-phenylfuran-2-carbaldehyde (0.43 g, 2.50 mmol) and *N*-methyl-1,2-phenylenediamine (0.5 mL, 4.4 mmol) were dissolved in a 1:1 mixture of ethanol and demineralized water (12 mL). Sodium bisulfite ($NaHSO_3$) (2.70 g, 26.0 mmol) was then added, and the mixture stirred at 70 °C overnight. On cooling the reaction mixture was diluted with water (25 mL) and extracted with ethyl acetate (3 x 25 mL), which was back washed with water (25 mL). The organic layers were separated, dried over anhydrous $Na_2SO_4$ and filtered. The filtrate was concentrated under reduced pressure, and the crude subjected to silica gel column chromatography (30% ethyl acetate in petroleum spirit) to afford B18 as an off-white solid (0.50 g, 73 %). $^{1}$H-NMR (400 MHz, DMSO-$d_6$): δ (ppm) 7.91-7.87 (m, 2H), 7.70-7.64 (m, 2H), 7.55-7.49 (m, 2H), 7.43-7.37 (m, 2H), 7.32-7.23 (m, 2H), 4.14 (s, 3H)

**4-(6-Chlorobenzofuran-2-yl)-2-(1*H*-pyrrol-2-yl)thiazole (B18-94)**

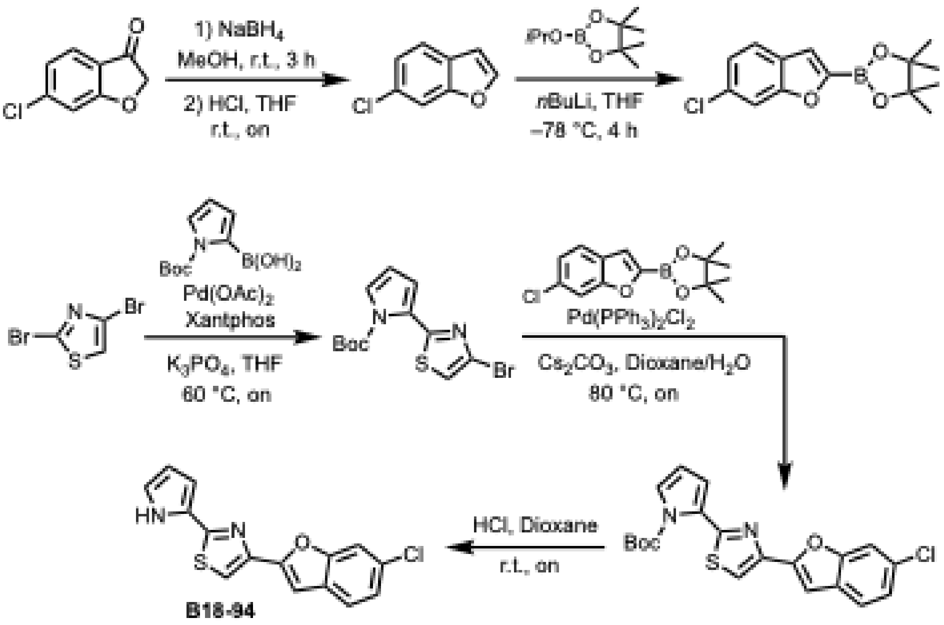


**6-Chlorobenzofuran**

Following the procedure of Zoubeidi *et al*[1]: 6-chlorobenzofuran-3(2H)-one (6.02 g, 35.71 mmol) was dissolved in MeOH (120 mL) to which sodium borohydride (3.64 g, 96.22 mmol) was slowly added portion wise at room temperature. The reaction mixture was stirred for 3 h before most of the MeOH was removed under reduced pressure. The mixture was dissolved in THF (30 mL), quenched with acetone (30 mL), and 5N HCl (60 mL) added. The reaction mixture was then stirred overnight which afforded a white precipitate. The resulting mixture was extracted with ethyl acetate (3 x 50 mL) and the organic layer washed with water (50 mL), dried over anhydrous $Na_2SO_4$, filtered and concentrated under reduced pressure. The crude was subjected to silica gel column chromatography (2% ethyl acetate in petroleum spirit) to afford 6-chlorobenzofuran as a colorless oil (4.93 g, 91%). Spectroscopic data were in accordance with literature[1].

$^1$H-NMR (400 MHz, $CDCl_3$): δ (ppm) 7.63-7.61 (d, 1H), 7.54-7.49 (m, 2H), 7.25-7.21 (dd, 1H), 6.77-6.73 (dd, 1H).

**2-(6-Chlorobenzofuran-2-yl)-4,4,5,5-tetramethyl-1,3,2-dioxaborolane**

Following a modified procedure of Zoubeidi *et al*[1]: 6-chlorobenzofuran (5.20 g, 34.08 mmol) was dissolved in anhydrous THF (114 mL) under an argon atmosphere, and the solution further purged with argon for 10 min before cooling to –78 °C. *n*BuLi (20 mL, 40 mmol, 2.0 M in hexanes) was then added dropwise, and the mixture stirred for 1.5 h before dropwise addition of 2-isopropoxy- 4,4,5,5-tetramethyl-1,3,2-dioxaborolane (15.21 mL, 74.60 mmol). After 2.5 h the reaction mixture was quenched with water (30 mL) and extracted with ethyl acetate (3 x 25 mL). The organic layers were washed with water (25 mL), dried over anhydrous $Na_2SO_4$ and filtered. The filtrate was concentrated under reduced pressure and the crude subjected to silica gel column chromatography (10-15 % ethyl acetate in petroleum spirit) to afford 2-(6-chlorobenzofuran-2-yl)-4,4,5,5-tetramethyl- 1,3,2-dioxaborolane as a slight orange colored oil (1.02 g, 11%). Spectroscopic data were in accordance with literature[1]. $^1$H-NMR (400 MHz, $CDCl_3$): δ (ppm) 7.56-7.51 (m, 2H), 7.36-7.35 (d, 1H), 7.24-7.20 (dd, 1H), 1.39 (s, 12H).

***tert*-Butyl 2-(4-bromothiazol-2-yl)-1*H*-pyrrole-1-carboxylate**

Following a modified procedure of Zoubeidi *et al*[1]: 2,4-dibromothiazole (0.52 g, 2.14 mmol), (1-(*tert*-butoxycarbonyl)-1*H*-pyrrol-2-yl) boronic acid (0.58 g, 2.75 mmol) and $K_3PO_4$ (0.96 g, 4.52 mmol, 2.1 eq) were dissolved in anhydrous THF (20 mL) and purged with argon for 20 min. Palladium acetate (58.5 mg, 0.26 mmol) and Xantphos (0.13 g, 0.22 mmol) were then added and the reaction mixture heated at 60 °C overnight under an argon atmosphere. On cooling to room temperature, the mixture was diluted with water (25 mL) and extracted with ethyl acetate (3 x 25 mL). The organic layers were washed with water (25 mL), dried over anhydrous $Na_2SO_4$ and filtered. The filtrate was concentrated under reduced pressure, and the crude subjected to silica gel column chromatography (2% ethyl acetate in petroleum spirit) to afford *tert*-butyl 2-(4-

bromothiazol-2-yl)-1*H*-pyrrole-1-carboxylate as a pale-yellow oil (0.29 g, 41%). $^{1}$H-NMR (400 MHz, $CDCl_3$): δ (ppm) 7.45-7.44 (dd, 1H), 7.29 (s, 1H), 6.70-6.67 (dd, 1H), 6.30-6.27 (t, 1H), 1.49 (s, 9H).

**_tert_-Butyl 2-(4-(6-chlorobenzofuran-2-yl)thiazol-2-yl)-1*H*-pyrrole-1-carboxylate**

Following a modified procedure of Zoubeidi *et al*[1]: *tert*-butyl 2-(4-bromothiazol-2-yl)-1*H*-pyrrole-1-carboxylate (78 mg, 0.24 mmol), 2-(6-chlorobenzofuran-2-yl)-4,4,5,5-tetramethyl-1,3,2-dioxaborolane (96 mg, 0.34 mmol) and $Cs_2CO_3$ (0.17 g, 0.52 mmol) were dissolved in a mixture of 1,4-dioxane (7 mL) and water (3mL). After purging the mixture with argon for 20 min, $PdCl_2(PPh_3)_2$ (35 mg, 0.05 mmol) was added, followed with further purging with argon for 10 min. The reaction mixture was then heated at 80 °C overnight under an argon atmosphere. On cooling to room temperature most of the 1,4-dioxane was removed under reduced pressure, and the mixture diluted with water (25 mL) and extracted with ethyl acetate (3 x 25 mL). The organic layers were washed with water (25 mL), dried over anhydrous $Na_2SO_4$ and filtered. The filtrate was concentrated under reduced pressure, and the crude subjected to silica gel column chromatography (5% ethyl acetate in petroleum spirit) to afford *tert*-butyl 2-(4-(6-chlorobenzofuran-2-yl)thiazol-2-yl)-1*H*-pyrrole-1-carboxylate as a pale-grey solid (41 mg, 43%). $^{1}$H-NMR (400 MHz, $CDCl_3$): δ (ppm) 7.67 (s, 1H), 7.51-7.48 (m, 2H), 7.24-7.21 (dd, 1H), 7.10 (s, 1H), 6.92-6.90 (d, 1H), 6.65-6.63 (d, 1H), 3.88 (s, 3H), 1.59 (s, 9H).

**4-(6-Chlorobenzofuran-2-yl)-2-(1*H*-pyrrol-2-yl)thiazole (B18-94)**

Following the procedure of Zoubeidi *et al*[1]: *tert*-butyl 2-(4-(6-chlorobenzofuran-2-yl)thiazol-2-yl)-1*H*-pyrrole-1-carboxylate (21.0 mg, 0.05 mmol) was dissolved in anhydrous 1,4-dioxane (2.2 mL) under an argon atmosphere. HCl (1.2 mL, 4M in dioxane) was then added dropwise, and the mixture stirred at room temperature overnight. The reaction mixture was placed under reduced pressure to remove the majority of the 1,4-dioxane, followed by dilution with water (25 mL) and extraction with ethyl acetate (3 x 25 mL). The organic layer was washed with water (25 mL), dried over anhydrous $Na_2SO_4$ and filtered. The filtrate was concentrated under reduced pressure, and the crude subjected to silica gel column chromatography (10% ethyl acetate in petroleum spirit) to afford 4-(6-chlorobenzofuran-2-yl)-2-(1H-pyrrol-2-yl)thiazole as a pale-grey solid (12 mg, 75%). Spectroscopic data were in accordance with literature[1]. $^1$H-NMR (400 MHz, $CDCl_3$): δ (ppm) 9.52-9.25 (bs, 1H), 7.58-7.44 (m, 3H), 7.25-7.21 (dd, 1H), 7.13 (s, 1H), 6.96-6.93 (m, 1H), 6.76-6.73 (m, 1H), 6.32-6.29 (m, 1H).

#### References


1. Zoubeidi, A.; Munuganti, R.S.N.; Bishop, J.L.; Thaper, D.; Vahid, S. Preparation of thiazole, benzothiazole derivatives and related heterocycles as transcription factor BRN2 inhibitors for the treatment of cancers and other diseases. WO2020069625A1. **2020**.
2. Jiang, Y.-q.; Jia, S.-h.; Li, X.-y.; Sun, Y.-m.; Li, W.; Zhang, W.-w. Xu, G.-q. An efficient $NaHSO_3$-promoted protocol for chemoselective synthesis of 2-substituted benzimidazoles in water. *Chem. Pap.* **2018**, *72*, 1265–1276.

#### $^{1}H$ NMR spectra

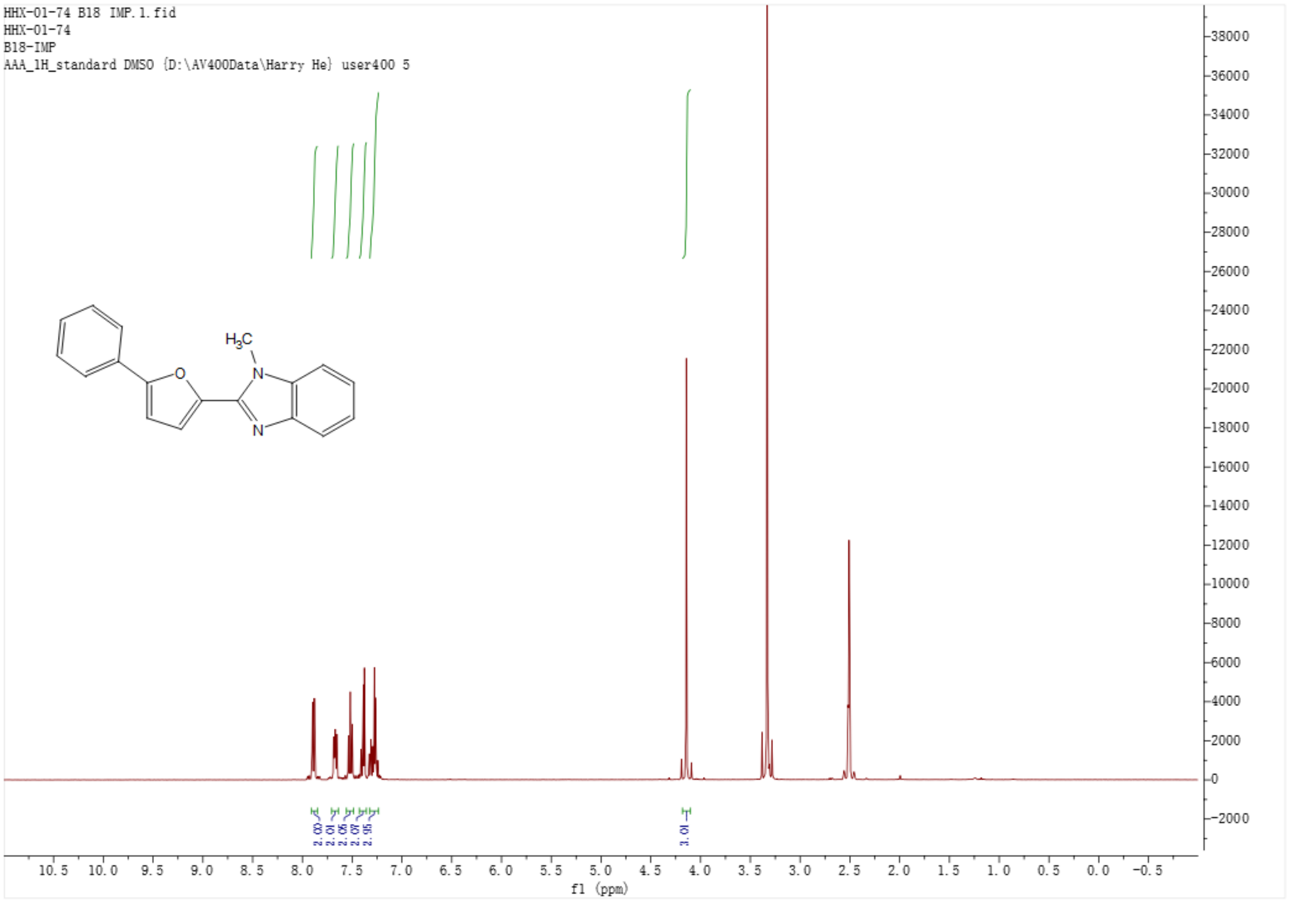


$^{1}H$ NMR spectrum of B18.

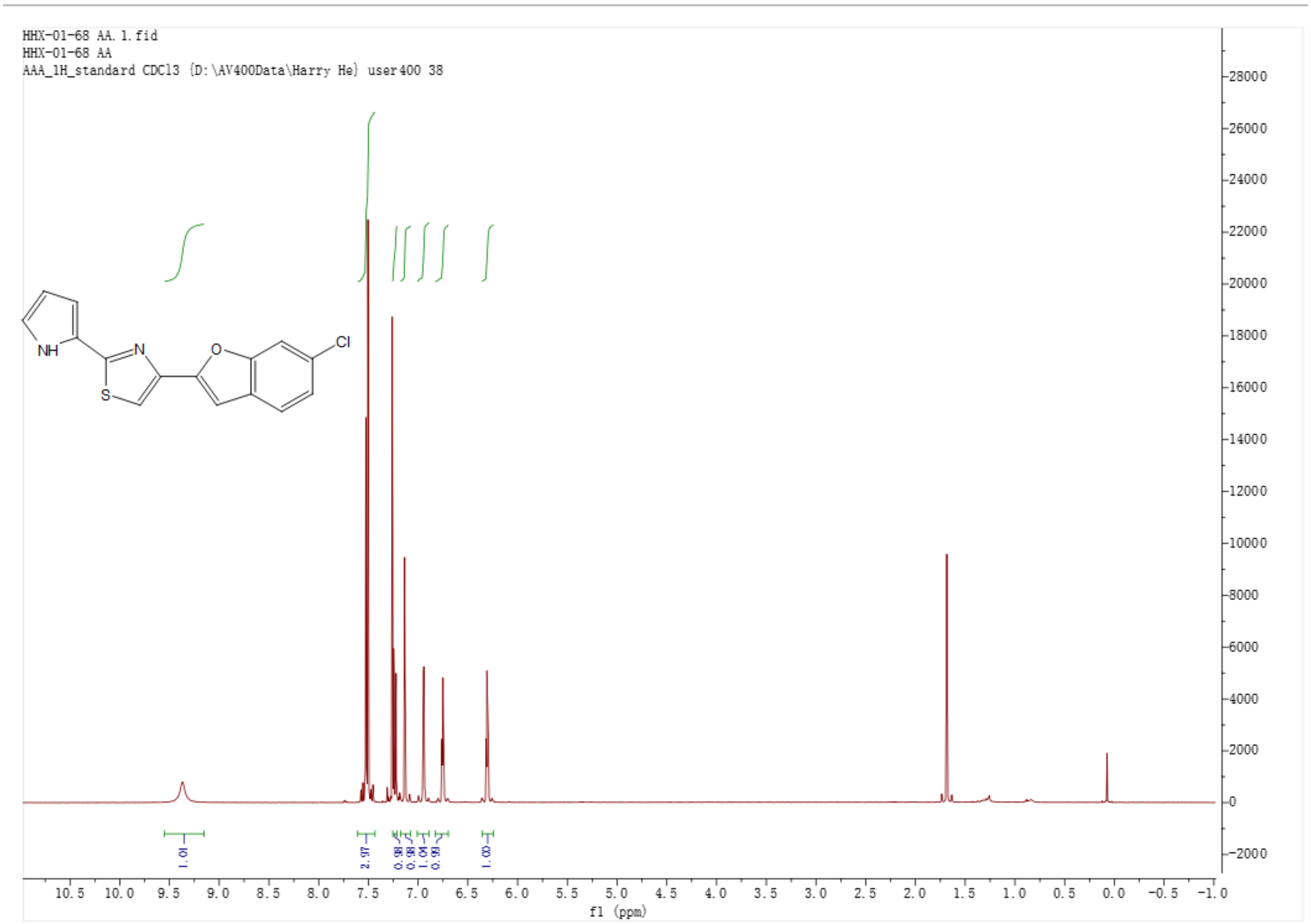


[1]H NMR spectrum of B18-94.

**X-Ray crystallography**

**B18** – Cambridge Structural Database number: 2451491
[https://www.ccdc.cam.ac.uk/solutions/software/csd/](https://www.ccdc.cam.ac.uk/solutions/software/csd/)

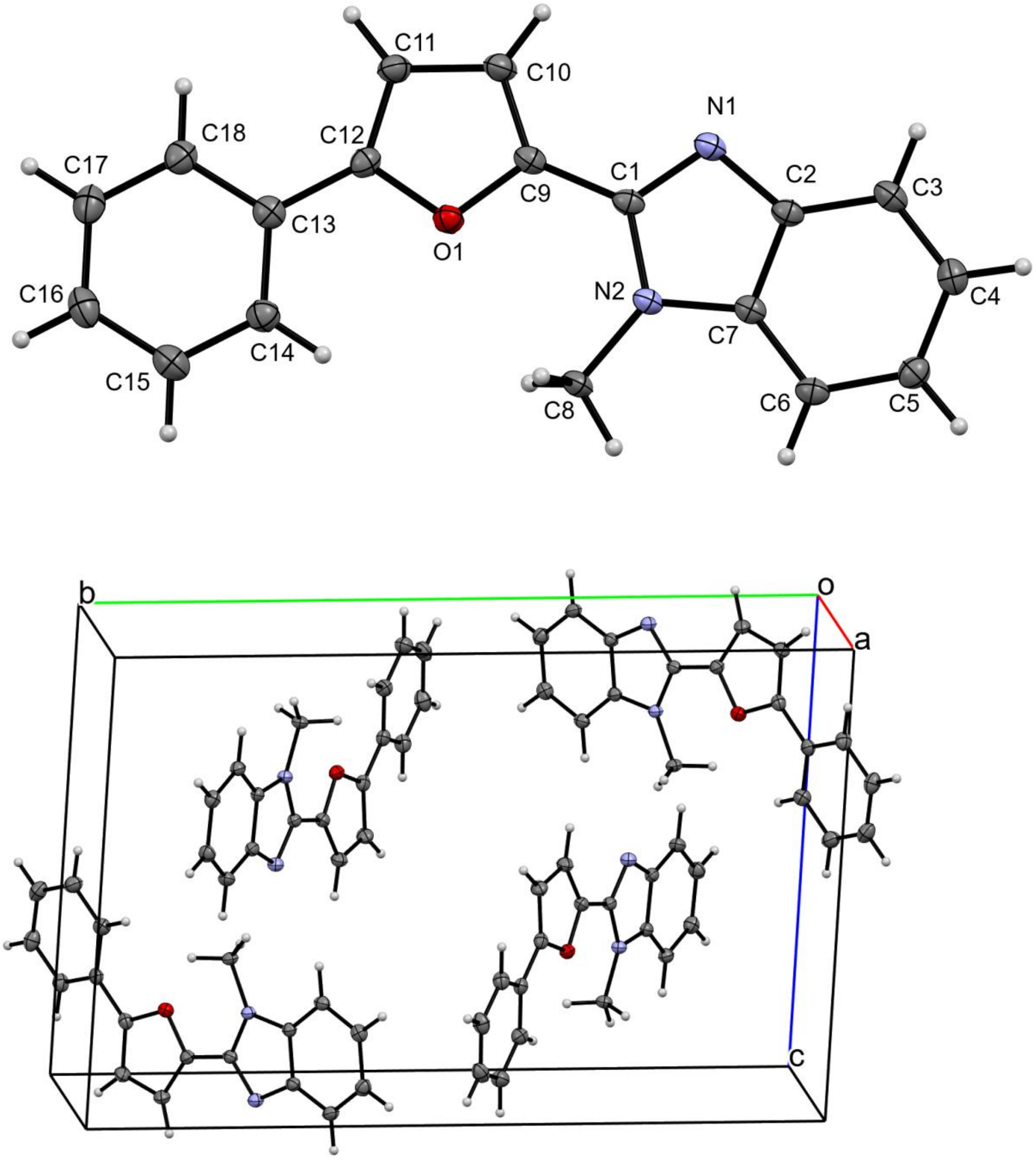


ORTEP diagram and unit cell for B18.

**B18-94** – Cambridge Structural Database number: 2451492
[https://www.ccdc.cam.ac.uk/solutions/software/csd/](https://www.ccdc.cam.ac.uk/solutions/software/csd/)

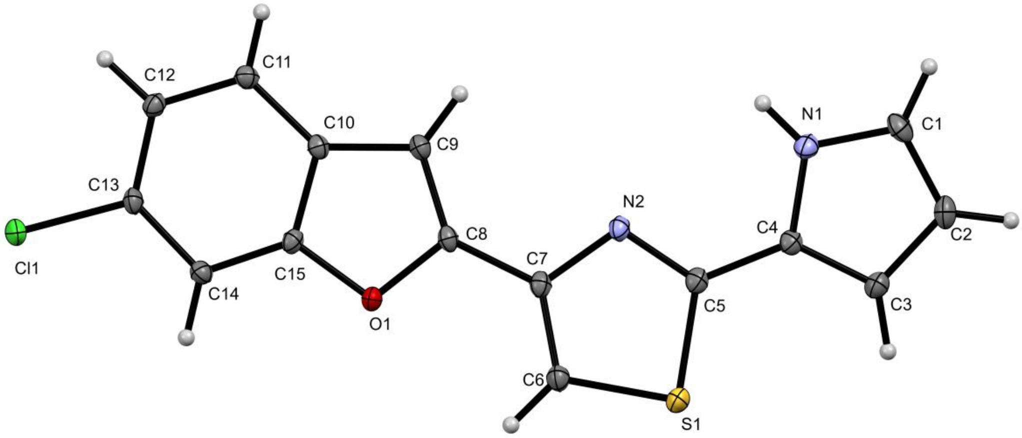


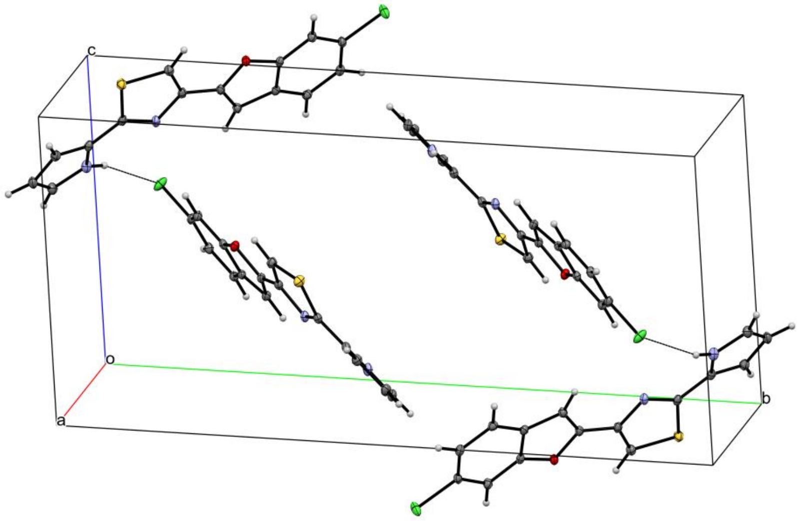


ORTEP diagram and unit cell for B18-94.
